# Modulation of multiple gene clusters expression by a single PAS-LuxR transcriptional regulator

**DOI:** 10.1101/2022.06.10.495441

**Authors:** Cláudia M. Vicente, Tamara D. Payero, Antonio Rodríguez-García, Eva G. Barreales, Antonio de Pedro, Fernando Santos-Beneit, Jesús F. Aparicio

## Abstract

PAS-LuxR transcriptional regulators are highly conserved enzymes governing polyene macrolide antifungal biosynthesis. PteF is one of such regulators, situated in the polyene macrolide filipin gene cluster from *Streptomyces avermitilis*. Its mutation leads to a drastic filipin production decline, but also to a severe loss of oligomycin production, an ATP-synthase inhibitor of macrolide structure, and a delay in sporulation, thus it has been considered as a transcriptional activator. Transcriptomic analyses were carried out in *S. avermitilis* Δ*pteF* and its parental strain *S. avermitilis* NRRL 8165 as control. Both strains were grown in YEME medium without sucrose, and samples were taken in the exponential and stationary growth phases. 257 genes showed altered expression in the PteF-deleted mutant, most of them in the exponential phase of growth. Surprisingly, despite PteF being an activator of filipin biosynthesis, a majority of the genes affected upon mutation showed overexpression thus suggesting a negative modulation of those genes. Genes affected were related to various metabolic processes, including genetic information processing; DNA, energy, carbohydrate, and lipid metabolism; morphological differentiation; and transcriptional regulation; among others, but particularly to secondary metabolite biosynthesis. Notably, ten secondary metabolite gene clusters out of 38 encoded by the genome, some of them encoding cryptic compounds, showed altered expression profiles in the mutant, suggesting a regulatory role for PteF wider than expected. Transcriptomic results were validated by quantitative reverse transcription polymerase chain reaction. These findings provide important clues to understand the intertwined regulatory machinery that modulates antibiotic biosynthesis in *Streptomyces*.

## INTRODUCTION

PAS-LuxR regulators are transcriptional factors that combine an N-terminal PAS sensory domain (Hefti et al., 2004) with a C-terminal helix-turn-helix (HTH) motif of the LuxR type for DNA-binding (Santos et al., 2012). The sensor domain is thought to detect a physical or chemical stimulus and regulate, in response, the activity of the effector domain (Möglich et al., 2009). The archetype of this class of regulators, PimM, was first identified in the antifungal pimaricin biosynthetic gene cluster from *Streptomyces natalensis* (Antón et al., 2007). It was characterised as a transcriptional activator of pimaricin biosynthesis since antifungal production was abolished upon gene deletion, and later its mode of action was characterised at the molecular level (Santos-Aberturas et al., 2011a). Since its discovery, homologous regulatory proteins have been found to be encoded in all known biosynthetic gene clusters of antifungal polyketides, and they have been shown to be functionally equivalent, to the extent that the production of pimaricin is restored in *S. natalensis ΔpimM* upon introduction of heterologous regulators of the PAS-LuxR class, such as *nysRIV* (nystatin), *amphRIV* (amphotericin), or *pteF* (filipin) into the strain (Santos-Aberturas et al. 2011b). Furthermore, introduction of a single copy of *pimM* into the amphotericin-producing strain *S. nodosus*, into the filipin-producing strain *S. avermitilis*, or into the rimocidin producing strain *S. rimosus*, boosted the production of all polyenes, thus indicating that these regulators are fully exchangeable (Santos-Aberturas et al. 2011b).

Although given their location in the chromosome, PAS-LuxR regulators were initially considered pathway-specific transcriptional regulators, recent results have shown that they should be considered as regulators with a wider range of implications. The canonical operator of PimM was used to search for putative targets of orthologous protein PteF in the genome of *S. avermitilis*, finding multiple binding sites located inside or upstream from genes involved in different aspects of both primary and secondary metabolism (Vicente et al,. 2015), thus suggesting that the regulator could govern those processes. Several of these operators were selected, and their binding to PimM DNA-binding domain demonstrated by electrophoretic mobility shift assays (EMSAs). As a proof of concept, the biosynthesis of the ATP-synthase inhibitor oligomycin whose gene cluster included two operators was studied (Vicente et al,. 2015). PteF deleted mutants, which showed a severe loss of filipin production and delayed spore formation in comparison to that of the wild-type strain (Vicente et al 2014), also showed a severe loss of oligomycin production and reduced expression of *olm* genes. Gene complementation of the mutant restored phenotype, thus demonstrating that PteF was able to co-regulate the biosynthesis of two related secondary metabolites, the polyketide macrolides filipin and oligomycin (Vicente et al., 2015). This cross-regulation could therefore be extended to all the clusters where operators were found, which suggests that PAS-LuxR regulators may affect a plethora of processes previously unforeseen. In this sense, the introduction of PAS-LuxR regulatory genes into different *Streptomyces* hosts has already proven useful for the awakening of dormant secondary metabolite biosynthetic genes (Olano et al., 2014; Martínez-Burgo et al., 2019).

Here we have used microarrays to study the transcriptome of *S. avermitilis* Δ*pteF* mutant in comparison with that of its parental strain in order to deepen our knowledge about the processes in which PteF is involved.

## MATERIALS AND METHODS

### Strains and cultivation

*S. avermitilis* NRRL 8165 and its mutant *S. avermitilis* Δ*pteF* (Vicente et al., 2014) were routinely grown in YEME medium (Kieser et al., 2000) without sucrose. Sporulation was achieved in TBO medium (Higgens et al. 1974) at 30°C.

### Nucleic acid extractions

RNA was extracted as described elsewhere (Vicente et al., 2014). Briefly, 2 ml from liquid cultures in YEME medium without sucrose were harvested by centrifugation and immediately frozen by immersion in liquid nitrogen. Cells were resuspended in lysis solution [600 μl RLT buffer (RNeasy mini kit; Qiagen); 6 μl 2-mercaptoethanol] and disrupted using sonicator (Ultrasonic processor XL; Misonix Inc.). RNeasy ® Mini kit (Qiagen) was used for RNA isolation using RNase-Free DNase Set (Qiagen) as specified by manufacturer, followed by two consecutive digestions with TURBO™ DNase from Ambion® according to the manufacturer’s instructions. Total RNA concentration was determined with a NanoDrop ND-1000 spectrophotometer (Thermo Scientific), and quality and integrity were checked in a Bioanalyzer 2100 apparatus (Agilent Technologies). Total genomic DNA (gDNA) was isolated from stationary phase cultures following the salting-out procedure (Kieser et al., 2000).

### Microarray hybridizations

The microarray experiment was performed using a common reference design (Gadgil et al. 2005). The microarray chip Custom Gene Expression Microarray, 8×15K (Agilent) was customized in order to include different sets of probes as indicated elsewhere (Beites et al., 2014). For each microarray hybridization, 10 pmol of Cy3-labelled cDNA obtained from total RNA were mixed with 80 pmol of Cy5-labelled genomic DNA as the common reference. Labelling, hybridization, washing and scanning conditions were carried out as indicated previously (Rodríguez-García et al. 2007; Guerra et al., 2012). Three biological replicates from independent cultures were made for each experimental condition.

### Identification of differentially transcribed genes

Microarray data were normalized and analysed with the Bioconductor package LIMMA (Linear Models for Microarray Analysis) (Smyth 2004; Smyth et al. 2005). Spot quality weights were estimated as indicated in the Supplementary section (Tables S1 and S2). Both local and global normalizations were used (Wu et al., 2005). Firstly, weighted medians of log_2_ Cy3/Cy5 intensities were calculated for print-tip correction and afterwards global Loess was applied (Smyth and Speed 2003). The normalized log_2_ of Cy3/Cy5 intensities is referred in this work as the Mg value, which is proportional to the abundance of transcripts for a particular gene (Mehra et al., 2006). The information from within-array spot duplicates (Smyth et al., 2005) and empirical array weights (Ritchie et al., 2006) were taken into account in the linear models (Smyth 2004). The Mg transcription values of the four experimental conditions were compared using two contrasts, mutant versus wild type, corresponding to the two studied growth phases (exponential and stationary). For each gene, the Mc value is the binary log of the differential transcription between the mutant and the wild strain. The Benjamini-Hochberg (BH) false-discovery rate correction was applied to the *p*-values. A positive Mc value indicates upregulation, and a negative one, downregulation. For each contrast a result was considered as statistically significant if the BH-corrected *p*-value was <0.05. In certain occasions, however, when the transcription profile of a gene matched that of genes statistically significant and functionally related, or for comparison with previous published results obtained by RT-qPCR or by EMSA assays (Vicente et al., 2014, 2015), we used an uncorrected *p*-value with a level of significance <0.05.

The microarray data are deposited in the National Center for Biotechnology Information-Gene Expression Omnibus under accession number GSE185887.

### Assessment of filipin and oligomycin production

Filipin production was quantified as described elsewhere (Barreales et al., 2020), whereas oligomycin was measured following the procedure described by Vicente et al (2015).

### Reverse transcription-quantitative PCR

Reverse transcription of total RNA was performed on selected samples with 5 μg of RNA and 12.5 ng/μl of random hexamer primer (Invitrogen) using SuperScript™ III reverse transcriptase (Invitrogen) as described previously (Barreales et al., 2018). Reactions were carried out on two biological replicates with three technical replicates each and appropriate controls were included to verify the absence of gDNA contamination in RNA and primer-dimer formation. Primers (see Table S3) were designed to generate PCR products between 97 and 153 bp, near the 5’ end of mRNA. The PCR reactions were initiated by incubating the sample at 95°C for 10 min followed by 40 cycles at 95°C for 15 s, 62-70°C (depending of the set of primers used) for 34 s, and 72 ºC for 30 s. To check the specificity of real-time PCR reactions, a DNA melting curve analysis was performed by holding the sample at 60°C for 60 s followed by slow ramping of the temperature to 95°C. Baseline and threshold values were determined by the StepOnePlus software. C_t_ values were normalized with respect to *rrnA1* mRNA (encoding 16S rRNA). Relative changes in gene expression were quantified using the Pfaffl method (2001) and the REST^©^ software (Pfaffl et al., 2002). The corresponding real-time PCR efficiency (E) of one cycle in the exponential phase was calculated according to the equation E = 10^[-1/slope]^ (Rasmussen, 2000) using 5-fold dilutions of genomic DNA ranging from 0.013 to 40 ng (n=5 or 6 with three replicates for each dilution) with a coefficient of determination *R*^2^ **>**0.99 (Fig. S1).

## RESULTS AND DISCUSSION

### Identification of genes with an altered expression profile in *S. avermitilis ΔteF* mutant

*S. avermitilis* Δ*pteF* and its parental strain *S. avermitilis* NRRL 8165 were grown in YEME medium without sucrose, and samples were taken at the end of the exponential and at the middle of the stationary growth phases (Fig. 1). Transcriptomic analysis was performed by microarray hybridization to assess the genes with an altered expression in the mutant when compared with the parental strain. Genomic DNA was used as a universal reference for all hybridizations. A result was considered as statistically significant if the BH-corrected *p*-value was <0.05. It is worth noting that these conditions are quite stringent, given that genes that constitute direct targets of PteF (e.g. the filipin polyketide synthases *pteA1* and *pteA2*; Vicente et al., 2014) are not statistically significant. With this criterion, microarrays analysis showed significant differences (with a fold change above or below +/-2) in the expression of 208 genes of the *pteF*-negative mutant at the end of the exponential phase, and 99 at the stationary phase of growth (Table 1; Fig. 2).

**Table 1:**
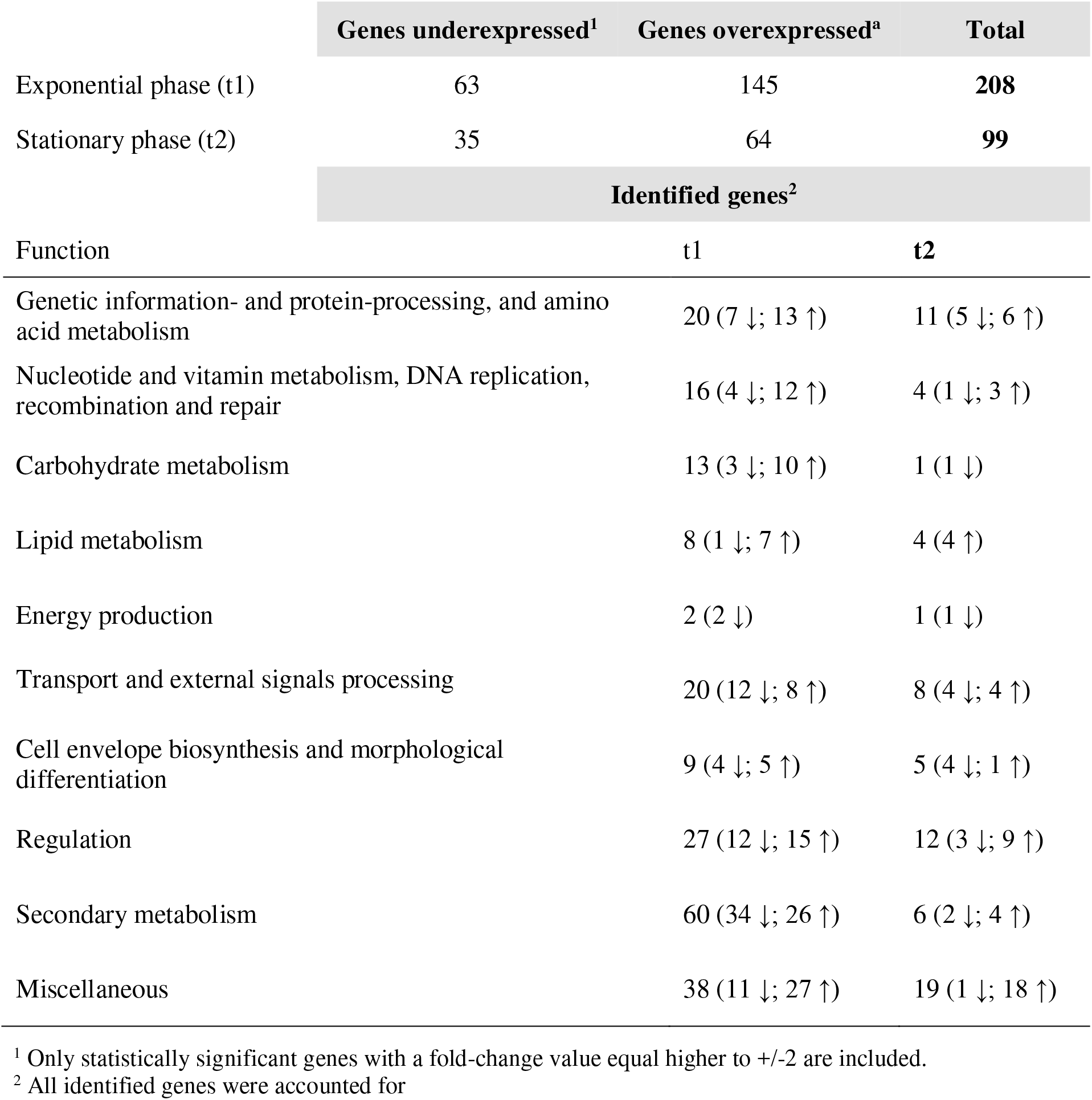
Differential transcription and functional classification of genes affected by *pteF* deletion.

**Fig. 1.**
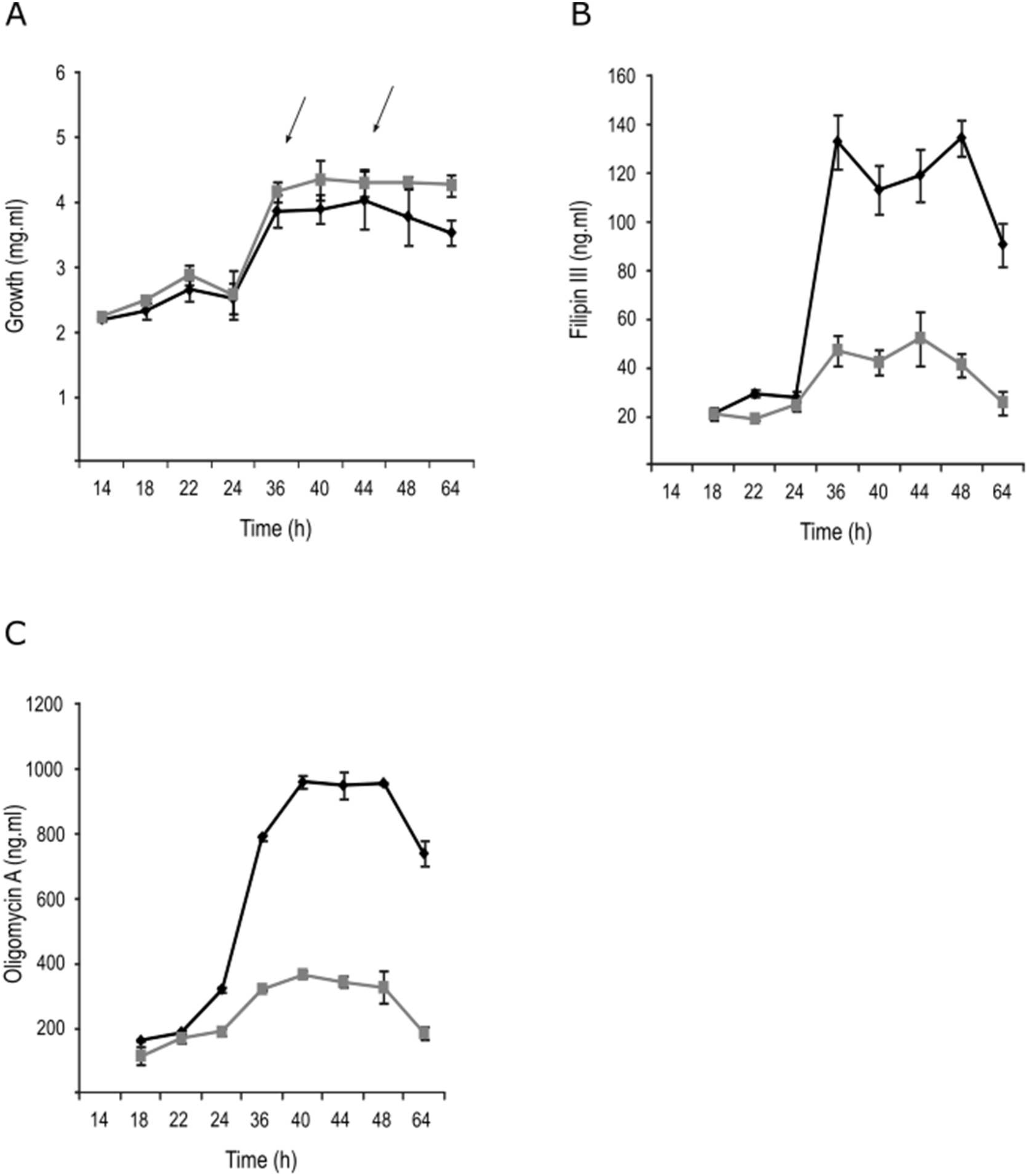
Growth and antibiotic production in YEME medium without sucrose. Strains *S. avermitilis* wt (black), and Δ*pteF* mutant (gray). A) Growth curves. B) Filipin production. (C) Oligomycin production. Arrows indicate RNA samples harvesting times.

**Fig. 2.**
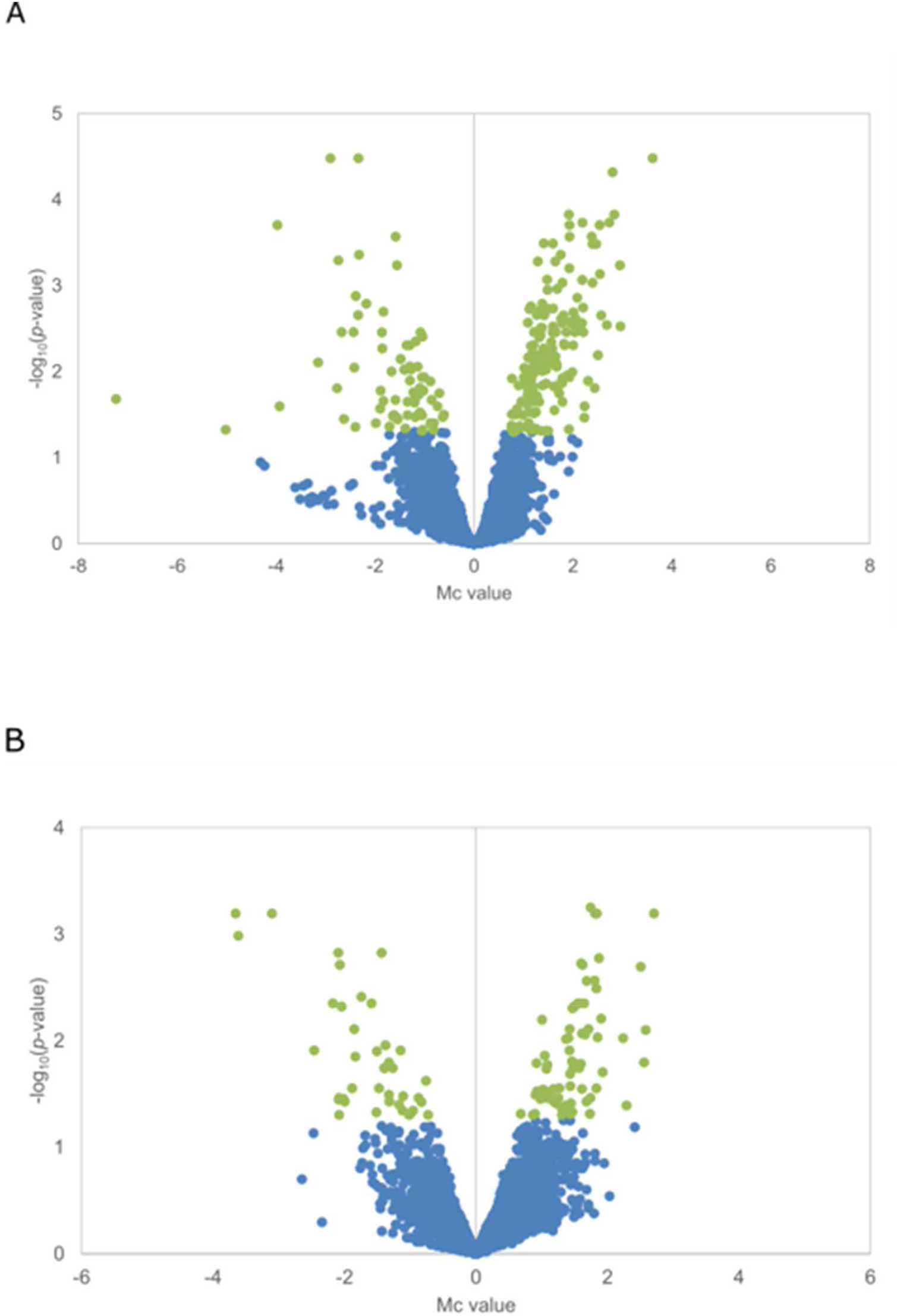
Differentially expressed genes in the mutant strain Δ*pteF*. Volcano plots show differential gene expression distribution during exponential phase (A) and stationary phase (B). Statistically significant genes are shown in green (log_10_ *p*-value ≥ 1.3).

Surprisingly, the lack of PteF resulted in the overexpression of a majority of the differentially transcribed genes, at both sampling times, thus indicating that this regulator acts as a negative modulator for those genes expression. This was unexpected given that PteF is an activator of both the antifungal filipin (Vicente et al., 2014) and the ATP-synthase inhibitor oligomycin (Vicente et al., 2015) biosynthesis.

These genes were related to different cellular processes, including genetic information processing; energy, carbohydrate, and lipid metabolism; DNA replication and repair; morphological differentiation; and transcriptional regulation, among others, but particularly to secondary metabolite biosynthesis (Table 1).

#### Genes involved in genetic information- and protein-processing, and amino acid metabolism

This group includes 24 genes that showed differential transcription in at least one of the sampling times (Table 1). These genes code for enzymes involved in amino acid metabolism (7 genes), proteins involved in transcription (8 genes, including 5 sigma factors), the ribosomal protein L28 (*SAV2675*), two putative acetyltransferases of ribosomal proteins (*SAV703* and *SAV758*), and enzymes involved in protein processing (5 genes) (Table S4).

Interestingly, while sigma factors *sig10* (*SAV898*), *sig13* (*SAV997*), and *sig60* (*SAV213*), and ribosomal proteins acetyltransferases *SAV703* and *SAV758* showed increased transcription levels in the mutant, *sig32* (*SAV3888*), *sig40* (*SAV4561*), the L28 ribosomal protein encoding gene *rpmB1*, and the *whiB*-like transcriptional factor *wblE* were clearly underexpressed in the mutant. The Wbl family of transcriptional factors is exclusive of actinobacteria and their members have been correlated with diverse roles in morphological differentiation and secondary metabolism (Fowler-Goldsworthy et al., 2011; Bush 2018).

Noteworthy, genes *rocA* (*SAV2723*) and *putA* (*SAV2724*), that encode delta-1-pyrroline-5-carboxylate dehydrogenase and proline dehydrogenase, respectively, and that have been related to proline catabolism (Menzel and Roth, 1981), and *rocD2* (*SAV7112*) and *SAV4551*, which encode putative ornithine aminotransferases, and are also involved in proline metabolism were underexpressed in the mutant, while *leuB* (*SAV2718*) which has been involved in valine, leucine, and isoleucine biosynthesis biosynthesis, *paaI* (*SAV1986*) that encodes a phenylacetic acid thioesterase, and putative cysteine desulfurase *SAV1061* were overexpressed.

#### Genes involved in nucleotide and vitamin metabolism, and DNA replication, recombination, and repair

Eighteen genes falling into this category were found to be differentially transcribed in the mutant (Table 1). Ten of them are involved in DNA replication, recombination and repair. Of these, seven putative transposases belonging to different families showed enhanced transcription in the mutant. Additionally, two genes involved in DNA repair, *ku2* (*SAV879*) which is probably involved in non-homologous DNA end-joining (Zhang et al., 2012), and *uvrD1* (*SAV3463*) that codes for a putative ATP-dependent helicase, were also upregulated. Conversely, *int12* (*SAV4626*), which encodes a tyrosine-family recombinase/integrase, showed reduced transcription levels at stationary phase.

The remaining genes were differentially transcribed only in the exponential phase. Four genes are involved in vitamin metabolism, three of them with lower transcription in the mutant, including cobalamin methylase *cobJ* (*SAV6407*) and adenosyltransferase *cobA* (*SAV6413*), and alkaline phosphatase *phoA* (*SAV5915*), which besides being part of the PhoRP two-component system (Sola-landa et al., 2008) is also involved in folate metabolism. The fourth gene, *thiC* (*SAV4265*) is a thiamine biosynthesis protein (Table S4). The rest of the genes are involved in purine metabolism, including *pgmA, purA*, and *purN*, all with enhanced transcription, and *cpdB*, with lower transcription.

#### Carbohydrate metabolism genes

Thirteen genes fall into this category, including four most likely belonging to the same operon (*SAV1009, galE5, mpg2, SAV1014*) and putatively involved in galactose metabolism, and showing enhanced transcription in the mutant. Other genes involved in the metabolism of this sugar were the alpha-galactosidase *agaB1* (*SAV1082*), which was underexpressed in the mutant, and the phosphoglucomutase *pgmA* (*SAV803*), which showed the opposite behavior. Interestingly, three genes of the tricarboxylic acid/glyoxylate cycle (citrate synthase *citA2*, citrate lyase *citE2*, and methylmalonyl-CoA mutase *meaA1*) were overexpressed in the mutant (Table S4).

#### Lipid metabolism genes

Nine genes related to lipid metabolism were differentially transcribed. These include the putative 3-oxoacyl-ACP synthase II *fabB2* (*SAV2944*), the acyl carrier protein *fabC4* (*SAV217*), the enoyl-CoA hydratase *echA1* (*SAV492*), and the acetyl/propionyl CoA carboxylase alpha subunit *accA2* (*SAV3866*), which are all presumably involved in fatty acid biosynthesis, and the 1-acylglycerol-3-phosphate O-acyltransferase *plsC1* (*SAV1485*) putatively involved in glycerophospholipid biosynthesis, among others. Interestingly, all these genes showed increased transcription in the mutant during the exponential phase except *fabB2* which was underexpressed (Table S4). However, during the stationary phase *fabB2* also showed enhanced transcription.

Noteworthy, the direct binding of the PteF orthologue PimM to the promoters of two of these genes was already demonstrated (Vicente et al., 2015), thus they have been included in Table S4 although they did not meet the statistical criteria. These were the acyltransferase *plsC1* (Yao and Rock 2013) whose transcription was increased in the mutant (Mc 0.88, uncorrected *p* value 0.0471), and *fabB2* whose transcription was reduced (Mc -0.84, uncorrected p-value 0.0410 in t1) or increased (Mc 1.12, *p*-value 0.0048 in t2) depending on the growth phase.

#### Energy production genes

Only three genes belonging to this group were found to be differentially transcribed in the mutant. All of them involved in oxidative phosphorylation and with reduced transcription in the mutant, two belonging to the operon *nuo* (*nuoJ1* and *nuoK1*), and the ATP synthase *atpF* (Table S4). Interestingly, all the genes belonging to the *nuo* operon (*SAV4837-SAV4850*), although in several cases not meeting the statistical criteria, showed the same decreased transcription profile in the mutant.

#### Transport and external signals processing

This group includes 25 genes that showed differential transcription in at least one of the sampling times (Table 1). Interestingly, twelve of them code or participate in the formation of ATP-binding cassette transporters (Table S4). Of these, four are putatively involved in sugar transport (*SAV1804, SAV2246, SAV2247*, and *SAV2609*) and showed reduced transcription in the mutant.

Four transporters belonging to the major facilitator superfamily showed differential transcription in the mutant, *SAV2455* with reduced transcription, and *SAV610*, the sulfate transporter *SAV4600, and SAV6941* with enhanced transcription.

Noteworthy, in agreement with the enhanced transcription of *SAV610*, the genes *fecC1* (*SAV600*) and *fecB* (*SAV602*) which constitute part of a putative ABC transporter iron(III)/siderophore transport system were also overexpressed. Based on protein similarity, SAV600-602 could constitute an ABC transport system homologous to the system FecBCD from *E. coli* involved in iron dicitrate transport (Staudenmaier et al., 1989). *SAV602* and *SAV610* genes flank a gene cluster involved in the biosynthesis of the siderophore nrp6 whose expression is also upregulated in the mutant (see below and Tables 2 and S4). Altogether these results suggest that the ABC system SAV600-602 and the transporter SAV610, would be involved in iron transport using the siderophore nrp6. These transcriptomic results are further supported by the direct binding of PimM to the promoters of *SAV602* and *SAV610* (Vicente et al., 2015).

**Table 2:**
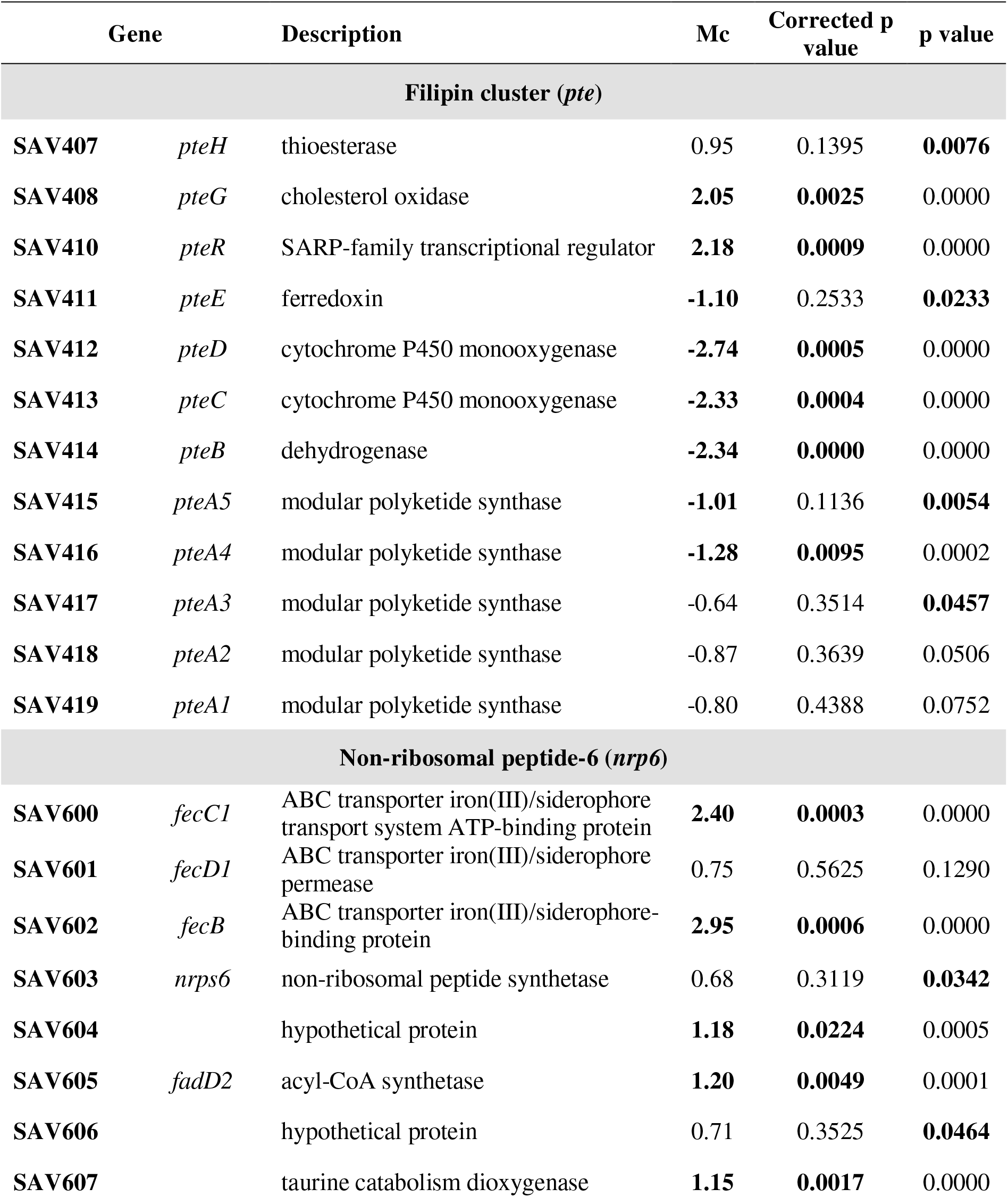
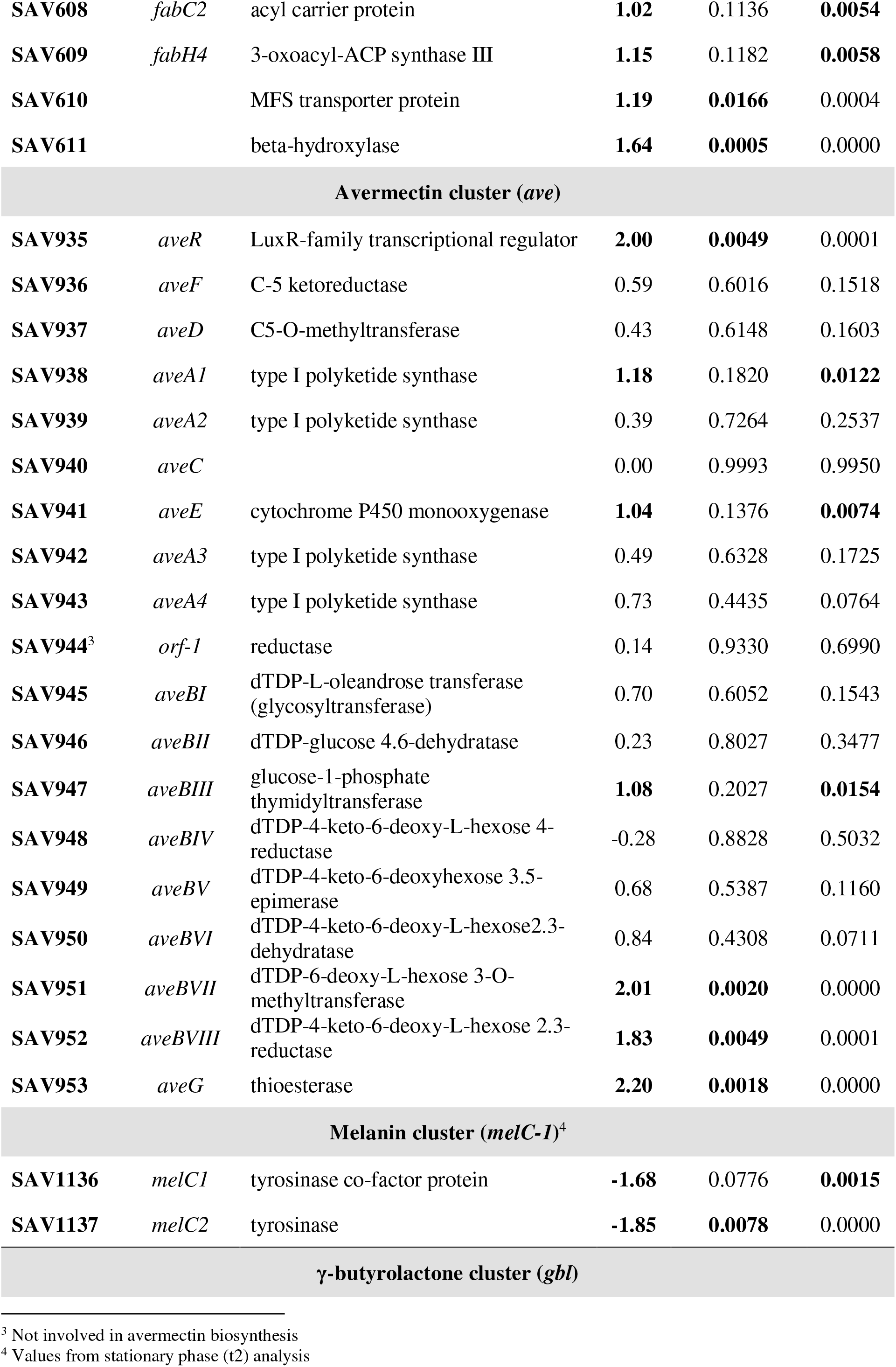
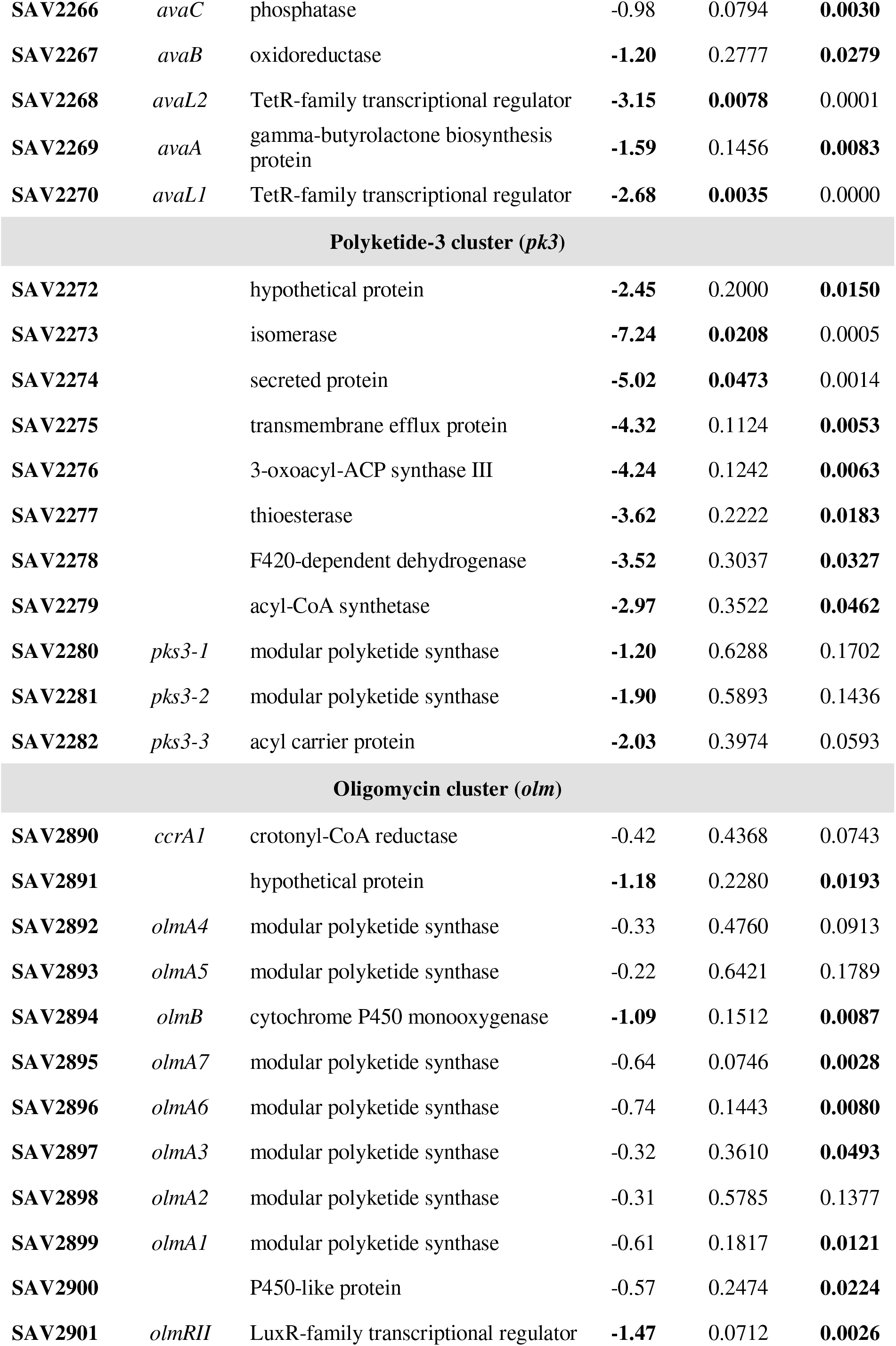
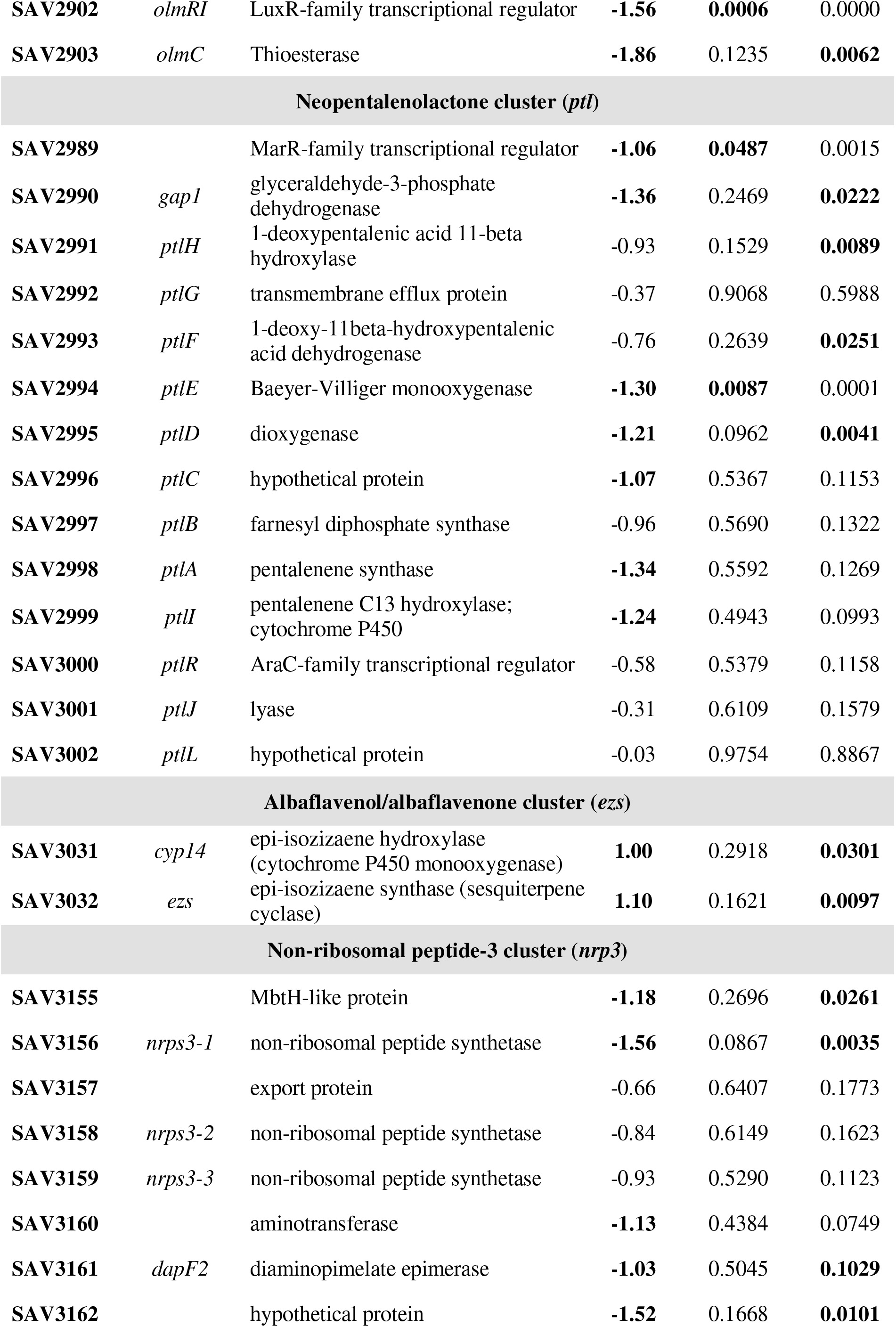
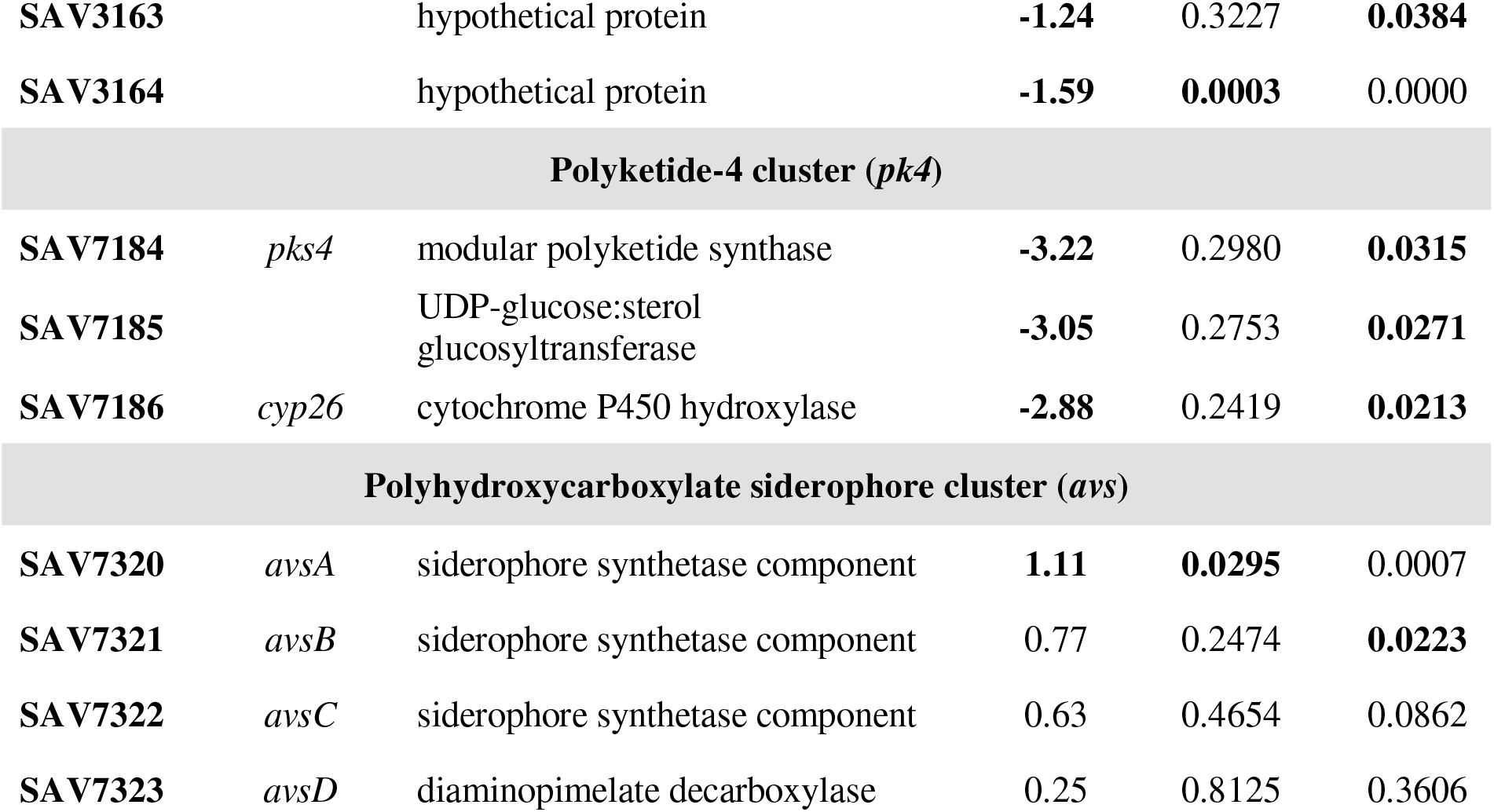
Transcriptional values of genes belonging to differentially expressed secondary metabolite gene clusters in *S. avermitilis ΔpteF* when compared to its parental strain. Differential transcription values *Mc* were obtained by subtracting mutant *Mg* values from parental strain *Mg* values. Only clusters whose transcription was affected by the mutation are included. The *p-*values are indicated in bold type when found statistically significant (see Materials and methods). For simplicity, designations ‘‘putative’’ have been removed.

#### Genes involved in cell envelope biosynthesis and morphological differentiation

This group includes eleven genes that showed differential transcription in at least one of the sampling times. These genes code for enzymes involved in cell envelope biosynthesis (the N-acetylmuramoyl-L-alanine amidase *ampD1*), and morphological differentiation (8 genes). The latter are particularly interesting because in *Streptomyces* morphological differentiation is usually accompanied by physiological differentiation (McCormick and Flärdh, 2012). The differential expression of genes involved in morphological differentiation was somehow expected given that *S. avermitilis ΔpteF* mutants show a delay in spore formation (Vicente et al., 2014).

Our results indicate that the transcriptional regulators *wlbE* and *bldC* that are associated to deficient phenotypes in spore formation (*white*) and in aerial mycelium development (*bald*), respectively, are underexpressed in the mutant. Similarly, the secreted subtilisin inhibitor *sit2* involved in morphological differentiation via *sigU* in *S. coelicolor* (Gordon et al., 2008), and *SAV2505* that encodes a DNA-binding protein orthologous to *S. lividans* transcriptional regulator ClgR which controls the expression of ATP-dependent protease Clp involved in morphological differentiation (Bellier et al., 2006), are also downregulated (Table S4). Interestingly, *clpC1* gene had also been proposed as direct PteF molecular target given PimM binding to its coding region (Vicente et al., 2015).

Conversely, the gene *ctpB* that encodes a cation-transporting P-type ATPase involved in *Bacillus subtilis* sporulation activation (Campo and Rudner, 2007), the gene *mreC* needed for spore cell-wall synthesis in *S. coelicolor* (Kleinschnitz et al., 2011), and both *kipI*, and its antagonist *kipA*, which have been involved in sporulation control in *B. subtilis* (Wang et al., 1997; Jacques et al., 2011), showed enhanced transcription in the mutant (Table S4).

#### Regulatory genes

As described here, a large set of genes with diverse functions are under the control of PteF, including several regulatory genes listed in the categories described above. This prompted us to analyze other possible transcriptional regulators differentially expressed in the mutants, as these could be mediators of the regulatory control. A complete list of regulatory genes whose expression is affected in the mutant is presented in Table S4.

A total of 31 transcriptional regulators showed a significant differential transcription in the mutant when compared with the parental strain. Such a large number reflects the pleiotropic nature of PAS-LuxR regulators (Vicente et al., 2014, 2015; Aparicio et al., 2016), and probably justifies all the biological processes affected by the mutation (see functional categories listed above).

Among the regulators controlled by PteF, is interesting to highlight eight directly involved in diverse secondary metabolites biosynthesis control, namely: *avaL2* (*SAV2268*) and *avaL1* (*SAV2270*), both TetR-family regulators putatively involved in the biosynthesis of a γ-butyrolactone (Ikeda et al., 2014); *avaR1* (*SAV3705*), which encodes the avenolide receptor protein (Kitani et al., 2011; Wang et al., 2014; Zhu et al., 2016); *olmRII* (*SAV2901*) and *olmRI* (*SAV2902*), both LuxR-family positive regulators of macrolide oligomycin biosynthesis (Yu et al., 2012); *pteR* (*SAV410*), the SARP-LAL regulator of the polyene macrolide filipin biosynthesis (Ikeda et al., 2014; Vicente et al., 2014; Payero et al., 2015); *aveR* (*SAV935*), a LAL-family positive regulator of avermectin biosynthesis (Kitani et al., 2009); and *SAV2989*, a MarR-family transcriptional regulator from the neopentalenolactone biosynthetic cluster (Ikeda et al., 2014). All these regulatory genes showed decreased transcription in the mutant, except for *pteR* and *aveR* that were overexpressed (Tables 2 and S4).

Interestingly, the expression of *olmRI* and *olmRII* genes had already been proven to be negatively affected by the lack of PteF (Vicente et al., 2015). Furthermore, *pteF*-deletion mutants showed a severe loss of oligomycin production, whereas gene complementation of the mutant restored parental-strain phenotype, and gene duplication in the wild-type strain boosted oligomycin production (Vicente et al., 2015). Similarly, *pteR* has also been reported as a PteF molecular target, via the action of another hierarchical regulator which would be activated by PteF (Vicente et al., 2014).

Besides the abovementioned regulators, it is also noteworthy the identification of *SAV2301*, that codes for a RedD orthologue, the transcriptional activator of the undecylprodigiosin pathway in *S. coelicolor* (Narva and Feitelson, 1990), *bldC* (*SAV4130*), a MerR-family regulator involved in morphological differentiation and secondary metabolite production in *S. coelicolor* (Hunt et al., 2005), and *cutS* (*SAV2404*), a sensor kinase involved in actinorhodin biosynthesis in *S. lividans* (Chang et al., 1996), all of them being down-regulated in the mutant (Table S4).

#### Secondary metabolite genes

The functional group more clearly affected by *pteF* deletion was that of genes involved in secondary metabolite biosynthesis (Table 1). In this category, when one or more genes critical for metabolite biosynthesis were found statistically significant, the transcription of other genes belonging to the same cluster with uncorrected p-values < 0.05 was also considered significant. Following this broader criterion, sixty one genes belonging to this group, regardless of regulatory genes mentioned above, showed a significant differential transcription in the mutant when compared with the parental strain in at least one of the sampling times (Table S4). Noteworthy, almost all genes were detected at the exponential growth-phase. In particular, those related to secondary metabolism precursor biosynthesis were only detected at this sampling time. These were: the ornithine aminotransferases *rocD3* (*SAV2285*) and *rocD2* (*SAV7112*), and the proline dehydrogenase *putA* (*SAV2724*), which were underexpressed; and the phosphoglucomutase *pgmA* (*SAV803*), the 3-isopropylmalate dehydrogenase *leuB* (*SAV2718*), the phosphoribosylglycinamide formyltransferase *purN* (*SAV3445*), and the putative citrate synthase *citA2* (*SAV3859*), which were overexpressed.

But the most striking results of microarray analyses were the identification of differential transcription in 67 genes (including regulatory genes) belonging to 10 out of the 38 putative secondary metabolite gene clusters encoded by *S. avermitilis* genome (Ikeda et al., 2014). Table 2 includes the transcriptional values of genes belonging to differentially expressed secondary metabolite gene clusters. For gene cluster boundaries definition we used StrepDB database (http://strepdb.streptomyces.org.uk) in conjunction with information described by Ikeda et al. (2014).

The secondary metabolites whose biosynthesis would be affected by *pteF* deletion were of different nature, and included: the polyketides filipin (*pte*), oligomycin (*olm*), avermectin (*ave*), and the product of *pks3*; the non-ribosomal peptides nrp3 and the siderophore nrp6; the vibrioferrin-like polyhydroxycarboxylate siderophore *avs*; the terpenoid neopentalenoketolactone (*ptl*); the γ-butyrolactone (*gbl*); and melanin (*melC-1*).

In all these clusters, the differential transcription of at least one key biosynthetic gene was observed. The number of genes affected were: eleven in the *nrp6* cluster (out of 12), ten (out of 13 and 14 respectively) in the case of the filipin and oligomycin clusters, eight (out of 11) in the case of the *pk3* cluster, seven in the case of the avermectin (out of 19) cluster, six in the *nrp3* cluster (out of 10), six in the *ptl* cluster (out of 14), five (out of 5) in the *gbl* cluster, and two in the *avs* (out of 4) and melanin *melC-1* (out of 2) clusters (Table 2).

Furthermore, a closer look at the transcription of the remaining genes of each of these clusters revealed that most of the genes of a given cluster, followed the same tendency. Figure 3 shows the transcription profiles of secondary metabolite gene clusters genes affected by the mutation including regulatory genes, and Table 2 the transcription values observed for each of the genes.

**Fig. 3.**
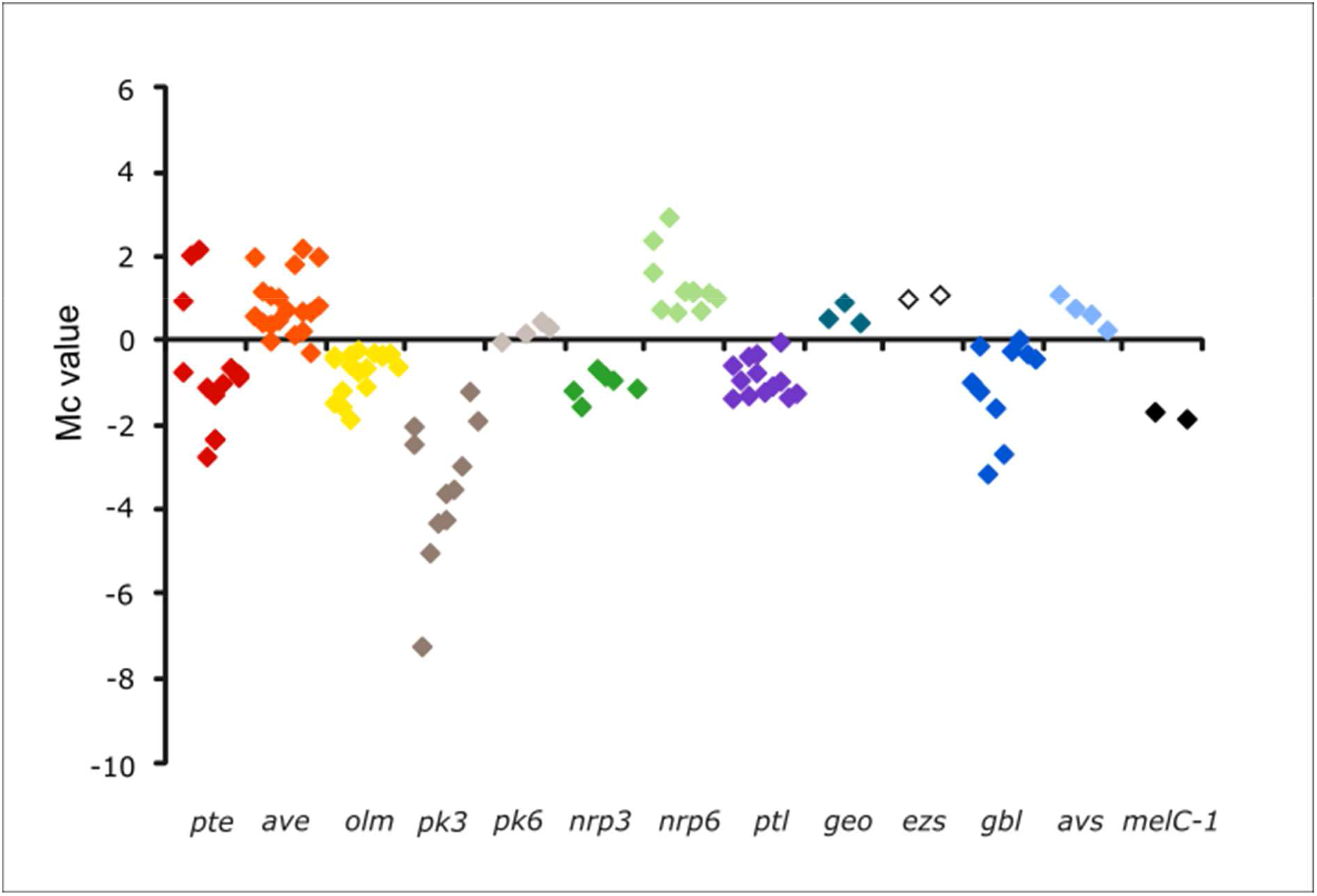
Transcription profiles of secondary metabolite gene clusters genes in *S. avermitilis ΔpteF*. Only clusters whose transcription was affected by the mutation are included. All the genes of a given cluster are shown in the plot, including regulatory genes. Coloured squares are the plots of differential transcription values for individual genes in the mutant. *pte*, filipin (red); *ave*, avermectin (orange); *olm*, oligomycin (yellow); *pk*, polyketide (gray); *nrp*, non-ribosomal peptide (green); *ptl*, neopentalenoketolactone (purple); *geo*, geosmin (teal); *ezs*, albaflavenol/albaflavenone (white); *gbl*, γ-butyrolactone (dark blue); *avs*, vibrioferrin-like siderophore (light blue); *melC*-1, melanin (black).

Seven of the secondary metabolite gene clusters showed an overall reduced transcription, including filipin *pte*, oligomycin *olm*, neopentalenoketolactone *ptl*, and melanin *melC*-1 clusters, the silent cluster for γ-butyrolactone *gbl*, and the cryptic gene clusters *pk3* and *nrp3*. On the opposite, three gene clusters showed overall enhanced transcription, including the macrolide avermectin *ave*, the siderophore *avs*, and the cryptic non-ribosomal peptide *nrp6* (Fig. 3).

Interestingly, besides the genes mentioned above, all the genes belonging to the clusters coding for the terpenoid albaflavenol/albaflavenone (*ezs*), and the cryptic polyketide *pk4* also followed the same tendency. In these cases, transcription values did not meet the statistical criteria, but their uncorrected *p*-values were < 0.05 in all instances (Table 2). In the case of the *ezs* genes (SAV3031-3032) they showed an average of two fold more transcription in the mutant, whereas *pk4* genes (*SAV7184-7186*) showed between 7 and 9 fold less transcription than in the parental strain.

### Filipin and Oligomycin production are strongly reduced in *S. avermitilis ΔpteF*

Although many of the metabolites whose biosynthesis would be affected by *pteF* deletion are of unknown structure (cryptic) and others are not produced under laboratory conditions (silent) (Ikeda et al., 2014), the production of two of them could be readily monitored in *S. avermitilis ΔpteF*. These were the antifungal pentaene filipin which is encoded by the *pte* cluster where the regulator is situated, and the ATP-synthase inhibitor oligomycin which is encoded by the *olm* cluster (Fig. 1). In both cases, production of the secondary metabolite was strongly reduced upon inactivation of the regulatory gene *pteF*. This is in agreement with the reduced transcription of most biosynthetic genes of both clusters (Fig. 3). The exceptions were the discrete thioesterase *pteH*, the cholesterol oxidase *pteG*, and the SARP-LAL regulator *pteR* of the filipin cluster, which were overexpressed. These results corroborate our previous observations by RT-qPCR (Vicente et al., 2014; Vicente et al., 2015).

### Validation of microarray results by using quantitative RT-PCR

Quantitative RT-PCR was used on reversed transcribed RNA samples to confirm that differential expression indicated by the microarray data was supported by an independent method. The selected genes covered a wide range of expression, including up-regulation and down-regulation. Twelve genes were validated including genes for the biosynthesis of filipin (*pteC, pteB, pteR, pteG*), oligomycin (*olmRI, olmRII, olmB*), avermectin (*aveR*), the isomerase of *pk3* cluster (*SAV2273*), one ABC transporter of the *nrp6* cluster (*fecB*), the alpha galactosidase *agaB1*, and the heat shock internal membrane protease *htpX1* (*SAV4891*).

Overall, the RT-qPCR data and microarray data showed a good concordance (Fig. S2). The range of dynamics for relative log_2_ fold change obtained from RT-qPCRs (−6.53 to +7.54) was higher than that obtained from Mc values from microarrays (−7.24 to +2.94), indicating that RT-qPCRs are more sensitive. This probably reflects on the Pearson’s correlation coefficient (*R*^2^) for the plot, resulting in a lower value than what could be expected. Nevertheless, the obtained value (*R*^2^ = 0.892) still indicates a good correlation of results.

### Concluding remarks

Up to date, PAS-LuxR regulator-encoding genes have been found only in polyene macrolide gene clusters, thus constituting a landmark of these type of clusters. In this context, they are transcriptional activators essential for the biosynthesis of the polyene encoded within the cluster. Their expression is a bottleneck in the biosynthesis of the antifungal and thus polyene production is easily incremented upon gene dosage increase (Aparicio et al., 2016). Additionally, heterologous gene complementation of mutants restores strain ability to produce the antifungal compound, thus proving that these regulators are highly conserved (Santos-Aberturas et al. 2011b). Recently, we have obtained evidence indicating that although these regulators were initially thought to be pathway-specific, they actually are regulatory proteins with a wider range of connotations in addition to polyene biosynthesis. Thus, PteF, the regulator of filipin biosynthesis, was proven to control oligomycin production in *S. avermitilis* (Vicente et al., 2015). This prompted us to propose that introduction of PAS-LuxR regulatory genes into *Streptomyces* species could prove useful for the awakening of dormant secondary metabolite biosynthetic genes (Vicente et al., 2014; 2015). This hypothesis was confirmed when PimM, the archetype of PAS-LuxR regulators was introduced into *S. albus* J1074, and production of the hybrid non ribosomal peptide-polyketide antimycin was activated (Olano et al., 2014). Recently, a similar result has been described in *S. albus* S4, where a PimM orthologue (the candicidin regulator FscRI) has been identified as required for antimycin production (McLean et al., 2016).

Here we have studied the transcriptome of *S. avermitilis* Δ*pteF* mutant in comparison with that of its parental strain. Our results corroborate our previous observations (Vicente et al., 2014, 2015), reinforcing the idea that PAS-LuxR regulators control many different cellular processes of bacterial metabolism at the transcriptional level, but in particular stress the importance of PAS-LuxR involvement on secondary metabolite biosynthesis.

Notably, ten (or twelve if we include *ezs* and *pk4* gene clusters) out of the 38 putative secondary metabolite gene clusters encoded by *S. avermitilis* genome (Ikeda et al., 2014) showed altered expression in the mutant. In some instances, the modified expression of biosynthetic genes of a given cluster could be explained by the effect of the mutation on the expression of one or more cluster-situated regulators. This is the case of the *aveR* regulator of the avermectin *ave* cluster, the regulators *avaL1* and *avaL2* of the γ-butyrolactone *gbl* cluster, the oligomycin regulators *olmRI* and *olmRII*, and the MarR regulator (*SAV2989*) of the pentalenolactone *ptl* cluster. AveR, the transcriptional activator of avermectin biosynthesis (Kitani et al., 2009), is overexpressed four-fold in the mutant and concomitantly the remaining genes of the *ave* cluster showed enhanced transcription. Conversely, OlmRI and OlmRI, positive regulators of oligomycin biosynthesis (Yu et al., 2012), showed decreased transcription in the mutant (Mc values -1.56 and -1.47 respectively), and so did the remaining genes of the cluster. It is not known whether AvaL1 and AvaL2 are positive regulators, but it is conceivable given that they show reduced transcription values upon mutation of the *pteF* gene (fold changes of 6.4 and 8.9, respectively) together with the remaining genes of the *gbl* cluster, including the γ-butyrolactone synthase *avaA*. Both AvaL1 and AvaL2 show convincing similarity to γ-butyrolactone receptor proteins, and although these proteins normally act repressing transcription of the synthase gene (Zou et al., 2014; Zhou et al., 2015; Barreales et al., 2020), there are cases that display the opposite behavior, like FarA from *S. lavendulae* that activates the transcription of the synthase *farX* (Kitani et al., 2010). The same occurs with the MarR regulator of the *ptl* cluster (Ikeda et al., 2014) whose transcription is diminished (2-fold) in the mutant as well as that of all *ptl* genes. In the remaining gene clusters there are no cluster-situated regulatory genes, thus the effect of the mutation must be explained either by a direct action of PteF on key biosynthetic genes or via the action of other regulatory genes. In this sense, 28 regulatory genes not situated in the clusters indicated above, most of them with unknown function, were differentially expressed upon mutation of *pteF* (Table S4).

To our knowledge, this is the second time a genome-wide transcriptomic study is conducted to describe the pleiotropic nature of a cluster-situated regulator, that of the regulator of lincomycin biosynthesis LmbU from *S. linconensis* (Li et al., 2020). Cross-regulation of disparate natural-product biosynthetic gene clusters by a cluster-situated regulator has already been described by several groups although not in genome-wide studies (Santamarta et al., 2011; Vicente et al., 2015; McLean et al., 2016). Moreover, the ability of some of these regulators to modulate the effects of regulators that act more globally (Huang et al., 2005), as well as the competition between global regulators (Santos-Beneit et al., 2009), have also been reported. Our findings go beyond, and indicate that PAS-LuxR regulators should be considered wide domain regulators. They affect the expression of multiple genes involved in both primary and secondary metabolism.

The findings reported here should provide important clues to understand the intertwined regulatory machinery that modulates antibiotic biosynthesis in *Streptomyces*, and suggest that the heterologous expression of PAS-LuxR regulators is likely to represent a powerful general strategy for novel bioactive natural product discovery.

## ACKNOWLEDGEMENTS

This work was supported by the Spanish Ministerio de Economía y Competitividad (Grant BIO2013-42983-P to JFA), F.P.U. fellowships of the Ministerio de Educación, Cultura y Deporte (AP2007-02055 to TDP, FPU13/01537 to AP), a contract from the Junta de Castilla y León co-financed by the European Social Fund (to EGB), and a fellowship from the Portuguese Fundação para a Ciência e a Tecnologia (SFRH/BD/64006/2009 to CMV).

