## Supplementary Material for "Modulation of multiple gene clusters expression by a single PAS-LuxR transcriptional regulator"

[b] *Institute of Biotechnology INBIOTEC, Parque Científico de León, Avda. Real, nº 1, 24006 León, Spain.*

[c] Present address: *TBI, CNRS, INRA, INSA, Université de Toulouse, Toulouse, France*

[d] Present address: *Institute of Sustainable Processes, University of Valladolid. Spain.*

### Supplementary Figures 1 and 2, Supplementary Tables S1-S4

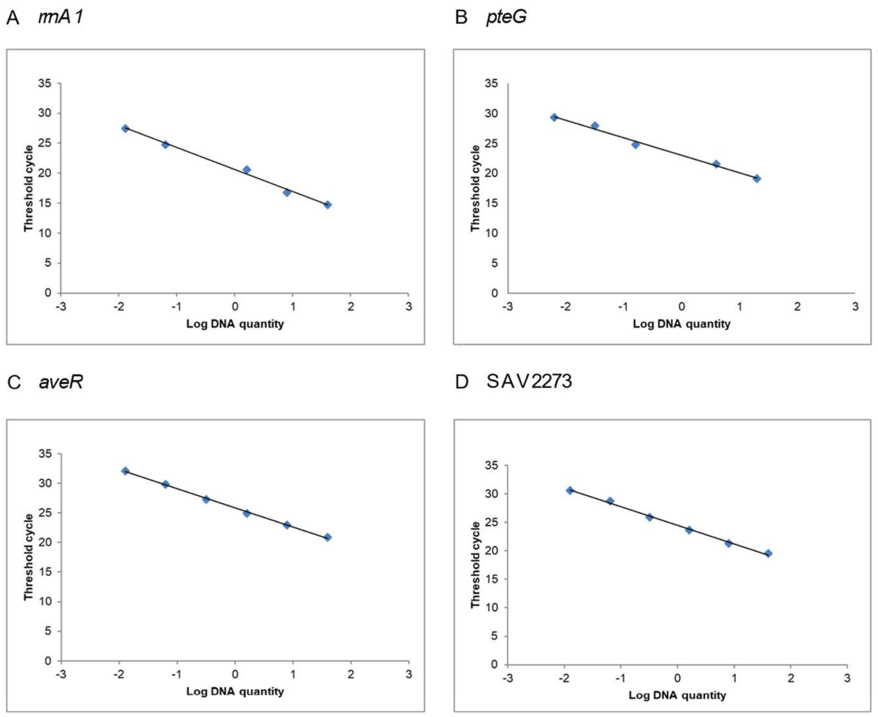


E

| | Slope | $R^2$ | Efficiency |
|---|---|---|---|
| rrnA1 fw/rev | -3,6512 | 0,992 | 0,88 |
| pteG fw/rev | -2.9279 | 0.991 | 1.20 |
| pteR fw/rev | -3.0556 | 0.989 | 1.12 |
| pteD fw/rev | -3,3057 | 0,991 | 1,01 |
| pteC fw/rev | -4,3989 | 0,998 | 0,69 |
| pteB fw/rev | -3,0027 | 0,997 | 1,15 |
| pteA4 fw/rev | -3,4593 | 0,977 | 0,95 |
| fecB fw/rev | -3,9301 | 0,996 | 0,80 |
| aveR fw/rev | -3,2291 | 0,998 | 1,04 |
| agaB1 fw/rev | -3,5398 | 0,994 | 0,92 |
| SAV2273 fw/rev | -3,2740 | 0,996 | 1,02 |
| olmRI fw/rev | -3,3578 | 0,996 | 0,99 |
| htpX1 fw/rev | -3,3973 | 0,995 | 0,97 |

**Fig. S1: Primer efficiency.** The efficiency of each set of primers was calculated according to the equation $E = 10^{[-1/slope]}-1$. Using 5-fold dilutions of genomic DNA, the resulting Ct values were plotted against the logarithm of the DNA quantity as shown in A (primers for *rrnA1*), B (primers for *pteG*), C (primers for *aveR*), or D (primers for SAV2273). Data are from three replicates, values represent the mean and the vertical bars ± SD. Panel E summarizes information obtained from all plotted data.

A

| Gene | Description | $log_2FC$ | Mc |
|---|---|---|---|
| *pteG* | cholesterol oxidase | 2.153 | 2.046 |
| *pteR* | SARP-LAL transcriptional regulator | 4.622 | 2.183 |
| *pteD* | cytochrome P450 monooxygenase | -4.655 | -2.744 |
| *pteC* | cytochrome P450 monooxygenase | -1.829 | -2.325 |
| *pteB* | dehydrogenase | -3.135 | -2.339 |
| *pteA4* | modular polyketide synthase | -0.544 | -1.278 |
| *fecB* | ABC transporter iron (III)/siderophore-binding protein | 7.537 | 2.946 |
| *aveR* | LuxR-family transcriptional regulator | 5.273 | 2.001 |
| *agaB1* | alpha-galactosidase | -1.969 | -2.907 |
| SAV2273 | isomerase | -6.532 | -7.240 |
| *olmRI* | LuxR-family transcriptional regulator | -1.278 | -1.560 |
| *htpX1* | heat shock protein, protease | 2.935 | 2.503 |

B

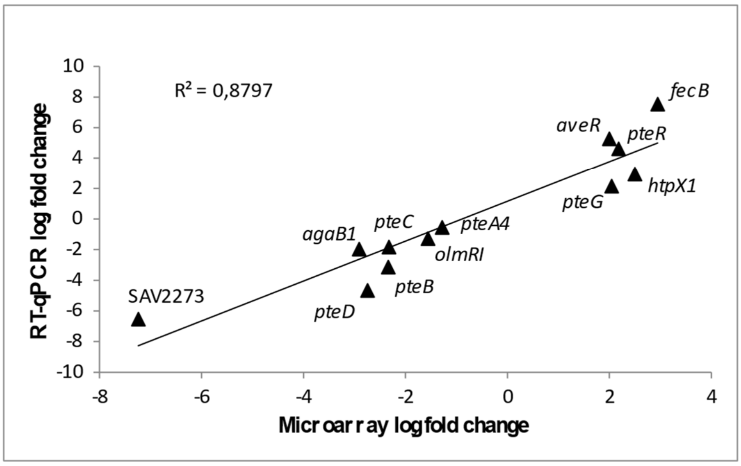


**Figure S2.- Validation of microarray results using RT-qPCR.** A) Comparison of the data obtained by RT-qPCR ($log_2FC$) and microarray analysis (Mc) for 12 different genes. B) Correlation between the results shown in panel A.

**Table S1: Determination of the quality flag for array spots.** The Feature Extraction software quantifies spot fluorescence and provides a set of Boolean values to assess the quality of the results. These quality indicators evaluate both red and green channel data of each spot (indicators are listed in the first row). Based on previous observations of the significance of each quality indicator, we classified the indicated combinations of Boolean values into a unique quality flag. Although spots with the best quality results were flagged as "1.00", other flag values were arbitrary.

| Step | gIsPosAndSignif | rIsPosAndSignif | gIsWellAboveBG | rIsWellAboveBG | gIsFeatNonUnifOL | rIsFeatNonUnifOL | gIsBGNonUnifOL | rIsBGNonUnifOL | gIsSaturated | rIsSaturated | **Flag** |
|---|---|---|---|---|---|---|---|---|---|---|---|
| 1st | 1 | 1 | 1 | 1 | 0 | 0 | 0 | 0 | 0 | 0 | 1.00 |
| 2nd | | 0 | | | | | | | | | 0.01 |
| 3rd | | | | | 0 | 0 | 0 | 0 | | | 0.95 |
| | | | | | 0<br>0<br>0<br>1 | 0<br>0<br>1<br>0 | 0<br>1<br>0<br>0 | 1<br>0<br>0<br>0 | | | 1.00 |
| | | | | | 0<br>1 | 1<br>0 | 1<br>0 | 0<br>1 | | | 0.85 |
| | | | | | 1 | 1 | 0 | 0 | | | 0.80 |
| | | | | | 1 | 0 | 1 | 0 | | | 0.70 |
| | | | | | 0 | 1 | 0 | 1 | | | 0.60 |
| | | | | | 0 | 0 | 1 | 1 | | | 0.50 |
| | | | | | 0<br>1<br>1<br>1 | 1<br>0<br>1<br>1 | 1<br>1<br>0<br>1 | 1<br>1<br>1<br>0 | | | 0.40 |
| | | | | | 1 | 1 | 1 | 1 | | | 0.30 |

**Table S2: Assigned weights to each spot flags.** To obtain spot quality weights that could be entered into the data analysis, we followed the idea behind the array weight estimation of Ritchie et al. (2006). In our case, the data of the 12 array hybridizations (4 conditions x 3 biological replicates) were normalized, the array weights calculated, and a linear model was fitted as indicated in Materials and Methods. In this initial stage, all spots were equally weighted irrespective of their quality flags. Spots with more reproducible transcription values —less variable— between replicates indicate a higher data quality. Hence, means of spot variances for each flag group were calculated. Spot weights were simply obtained by the normalized inverse of the mean variance.

| **Quality flag** | **Mean of variances** | **Quality weight** | **Percentage of total spots** |
|---|---|---|---|
| 1.00 | 4.296 | 1.000 | 62.601% |
| 0.80 | 4.827 | 0.890 | 4.110% |
| 0.95 | 5.209 | 0.825 | 0.015% |
| 0.01 | 6.366 | 0.675 | 32.611% |
| 0.60 | 11.985 | 0.358 | 0.005% |
| 0.40 | 20.589 | 0.209 | 0.002% |
| 0.85 | 23.558 | 0.182 | 0.002% |
| 0.30 | 35.496 | 0.121 | 0.051% |
| 0.70 | 36.000 | 0.119 | 0.006% |
| 0.50 | 44.424 | 0.097 | 0.597% |

**Table S3: Sequence of primers used for qPCR.**

| Primer | Sequence (5'to 3') | Transcripts quantification | Product size (bp) |
|---|---|---|---|
| Q-rrnA1 fw<br>Q-rrnA1 rev | GACGCAACGCGAAGAACC<br>TGCGGGACTTAACCCAACATC | *rrnA1* | 137 |
| Q-pteG fw<br>Q-pteG rev | CGGTGCCGACCACGAC<br>GCGGGGCAGACTACGATCAC | *pteG* | 100 |
| Q-pteR fw<br>Q-pteR rev | GGCCCTGCTGCTCATCC<br>CCTGTTCCCGCCTTTGTG | *pteR* | 153 |
| Q-pteD fw<br>Q-pteD rev | GCCAGCATGTCGTCCAAGAG<br>CTTCCCCCTGATCGGTGTC | *pteD* | 141 |
| Q-pteC fw<br>Q-pteC rev | GTGCTCCGGCTCGTCCTG<br>CCTGCTGCGCGACTCCTC | *pteC* | 118 |
| Q-pteB fw<br>Q-pteB rev | CCAGCGAGGACCACACG<br>GCCGAAGCAGGCGTTCC | *pteB* | 116 |
| Q-pteA4 fw<br>Q-pteA4 rev | TCAGGCCAAGGAAGTACGAGAC<br>CGCCATGTGGGACGACTAC | *pteA4* | 125 |
| Q-sav602 fw<br>Q-sav602 rev | CGGGCGCCTTGGTGAAC<br>GCCACCGGCGACAAGGC | *fecB* | 97 |
| Q-sav935 fw<br>Q-sav935 rev | GCACGGTGAAACTGCTCGTC<br>GTGGTCCGCGGGAAGCC | *aveR* | 131 |
| Q-sav1082 fw<br>Q-sav1082 rev | GCAGACCGAGTGTGGGAGAG<br>CCGGTGGGCTGCCATTCG | *agaB1* | 115 |
| Q-sav2273 fw<br>Q-sav2273 rev | GCCAGCGGATCGATGTACGG<br>AGTACCTCGCCCTGTGGAAC | *SAV2273* | 97 |
| Q-sav2902 fw<br>Q-sav2902 rev | GAGGTCGTCCGGGAAAGGAG<br>TCGGCCTGATCGCTCTGC | *olmRI* | 104 |
| Q-sav4891 fw<br>Q-sav4891 rev | GTCGTCGCACTCTTCATCGC<br>GGCCTCGAACTCGCTCACC | *htpX1* | 108 |

**Table S4: Differentially expressed genes in *S. avermitilis ΔpteF* when compared to its parental strain**. Genes are ordered firstly by functional class, secondly by reduced or enhanced transcription, and then by chromosomal position with the aim of highlighting the coincidence of profiles among clustered genes. The primary annotation source is the StrepDB server (http://strepdb.streptomyces.org.uk). Some genes are included in more than one functional category because of they are implicated in several processes. The *p*-values are indicated in bold type when found statistically significant (see Materials and methods). A few genes that did not meet criteria are also included (see footnotes). For simplicity, designations ''putative'' have been removed.

| Gene | | Description | Mc | Corrected p value | p value |
|---|---|---|---|---|---|
| **Genes involved in genetic information- and protein-processing, and amino acid metabolism** | | | | | |
| **t1** | | | | | |
| **SAV2723** | *rocA* | delta-1-pyrroline-5-carboxylate dehydrogenase | **-1.83** | **0.0219** | 0.0005 |
| **SAV2724** | *putA* | proline dehydrogenase | **-1.41** | **0.0095** | 0.0002 |
| **SAV2795** | *zmp4* | griselysin (secreted neutral zinc metalloprotease) | **-1.04** | **0.0473** | 0.0014 |
| **SAV4551** | | ornithine aminotransferase | **-1.89** | **0.0166** | 0.0004 |
| **SAV4561**[i] | *sig40* | RNA polymerase ECF-subfamily sigma factor | -0.81 | 0.1009 | **0.0045** |
| **SAV4562**[a] | | hypothetical protein | **-1.04** | **0.0039** | 0.0000 |
| **SAV7112** | *rocD2* | ornithine aminotransferase | **-1.59** | **0.0212** | 0.0005 |
| **SAV213** | *sig60* | RNA polymerase ECF-subfamily sigma factor | **1.78** | **0.0135** | 0.0003 |
| **SAV321** | *prpB1* | magnesium or manganese-dependent protein phosphatase | **1.58** | **0.0054** | 0.0001 |
| **SAV459** | *hsp18_1* | heat shock protein | **1.20** | **0.0106** | 0.0002 |
| **SAV692** | *hsp18_2* | heat shock protein | **1.93** | **0.0002** | 0.0000 |
| **SAV703** | | acetyltransferase | **1.27** | **0.0022** | 0.0000 |
| **SAV758** | | acetyltransferase | **1.16** | **0.0063** | 0.0001 |
| **SAV826** | *cspD1* | cold-shock protein | **1.48** | **0.0011** | 0.0000 |
| **SAV898** | *sig10* | RNA polymerase ECF-subfamily sigma factor | **2.54** | **0.0007** | 0.0000 |
| **SAV997** | *sig13* | RNA polymerase ECF-subfamily sigma factor | **1.42** | **0.0071** | 0.0001 |
| **SAV1061** | | cysteine desulfurase | **2.74** | **0.0002** | 0.0000 |
| **SAV1986** | *paaI* | phenylacetic acid degradation protein | **1.37** | **0.0031** | 0.0000 |

| | | | | | |
|---|---|---|---|---|---|
| **SAV2718** | *leuB* | 3-isopropylmalate dehydrogenase | **2.23** | **0.0252** | 0.0006 |
| **SAV4891** | *htpX1* | heat shock protein, protease | **2.50** | **0.0064** | 0.0001 |
| **t2** | | | | | |
| **SAV2675** | *rpmB1* | ribosomal protein L28 | **-1.17** | **0.0397** | 0.0005 |
| **SAV3016** | *wblE* | WhiB-family transcriptional regulator | **-1.12** | **0.0451** | 0.0006 |
| **SAV3888** | *sig32* | RNA polymerase ECF-subfamily sigma factor | **-3.61** | **0.0010** | 0.0000 |
| **SAV4561**[a] | *sig40* | RNA polymerase ECF-subfamily sigma factor | -0.75 | 0.1641 | **0.0074** |
| **SAV4820** | *cspD6* | cold-shock protein | **-2.02** | **0.0350** | 0.0004 |
| **SAV459** | *hsp18_1* | heat shock protein | **1.69** | **0.0027** | 0.0000 |
| **SAV692** | *hsp18_2* | heat shock protein | **1.84** | **0.0006** | 0.0000 |
| **SAV758** | | acetyltransferase | **1.02** | **0.0286** | 0.0003 |
| **SAV826** | *cspD1* | cold-shock protein | **1.09** | **0.0168** | 0.0001 |
| **SAV898** | *sig10* | RNA polymerase ECF-subfamily sigma factor | **1.61** | **0.0283** | 0.0003 |
| **SAV1061** | | aminotransferase | **1.67** | **0.0087** | 0.0000 |
| **Genes involved in nucleotide and vitamin metabolism, and DNA replication, recombination and repair** | | | | | |
| **t1** | | | | | |
| **SAV5915** | *phoA* | alkaline phosphatase | **-2.39** | **0.0013** | 0.0000 |
| **SAV6215** | *cpdB* | 2',3'-cyclic-nucleotide 2'-phosphodiesterase | **-1.57** | **0.0341** | 0.0009 |
| **SAV6407** | *cobJ* | precorrin methylase | **-1.03** | **0.0166** | 0.0004 |
| **SAV6413** | *cobA* | cob(I)alamin adenolsyltransferase | -0.61 | **0.0315** | 0.0008 |
| **SAV47** | | IS605 family IS1136A-like transposase | **1.75** | **0.0135** | 0.0003 |
| **SAV61** | | IS605 family IS1136A0-like transposase | **1.71** | **0.0145** | 0.0003 |
| **SAV120** | | IS701 family ISRhosp2-like transposase | **1.25** | **0.0293** | 0.0007 |
| **SAV288** | | IS630 family ISMac13-like transposase | **1.34** | **0.0082** | 0.0001 |
| **SAV757** | | IS630 family IS885-like transposase | **1.45** | **0.0095** | 0.0002 |
| **SAV803** | *pgmA* | phosphoglucomutase | **1.67** | **0.0155** | 0.0003 |
| **SAV821** | | IS200/IS605 family ISFsp4-like transposase | **1.60** | **0.0003** | 0.0000 |
| **SAV879** | *ku2* | Ku70/Ku80 protein | **1.01** | **0.0135** | 0.0003 |
| **SAV3445** | *purN* | phosphoribosylglycinamide formyltransferase | **1.29** | **0.0005** | 0.0000 |

| | | | | | |
|---|---|---|---|---|---|
| **SAV3463** | *uvrD1* | ATP-dependent DNA helicase | 0.77 | **0.0312** | 0.0008 |
| **SAV4139** | *purQ* | phosphoribosyl formylglycinamidine synthase I | 0.85 | **0.0313** | 0.0008 |
| **SAV4265** | *thiC* | thiamine biosynthesis protein | **1.60** | **0.0018** | 0.0000 |
| | | **t2** | | | |
| **SAV4626** | *int12* | tyrosine-family recombinase/integrase | **-1.51** | **0.0125** | 0.0001 |
| **SAV288** | | IS630 family ISMac13-like transposase | **1.17** | **0.0358** | 0.0004 |
| **SAV289** | | IS701 family ISAzvi8-like transposase | **1.43** | **0.0078** | 0.0001 |
| **SAV757** | | IS630 family IS885-like transposase | **1.51** | **0.0160** | 0.0001 |
| | | **Carbohydrate metabolism genes** | | | |
| | | **t1** | | | |
| **SAV1082** | *agaB1* | alpha-galactosidase | **-2.91** | **0.0000** | 0.0000 |
| **SAV2060** | | polysaccharide deacetylase, secreted | **-1.06** | **0.0116** | 0.0002 |
| **SAV6534** | | glycosyltransferase | **-1.30** | **0.0126** | 0.0002 |
| **SAV649** | *ard* | phosphotransferase | **1.47** | **0.0009** | 0.0000 |
| **SAV803** | *pgmA* | phosphoglucomutase | **1.67** | **0.0155** | 0.0003 |
| **SAV1009** | | glycosyltransferase | **1.55** | **0.0067** | 0.0001 |
| **SAV1011** | *galE5* | UDP-glucose 4-epimerase | **1.94** | **0.0027** | 0.0000 |
| **SAV1013** | *mpg2* | mannose-1-phosphate guanyltransferase | **1.99** | **0.0101** | 0.0002 |
| **SAV1014** | | NDP-hexose 4-ketoreductase | **1.47** | **0.0077** | 0.0001 |
| **SAV1911** | *ccrA2* | crotonyl-CoA reductase | **1.32** | **0.0040** | 0.0000 |
| **SAV1912** | *meaA1* | methylmalonyl-CoA mutase, coenzyme B12-dependent α-subunit | **1.93** | **0.0003** | 0.0000 |
| **SAV1913** | *citE2* | citrate lyase beta chain | **1.37** | **0.0016** | 0.0000 |
| **SAV3859** | *citA2* | citrate synthase | **1.38** | **0.0144** | 0.0003 |
| | | **t2** | | | |
| **SAV1082** | *agaB1* | alpha-galactosidase | **-1.26** | **0.0180** | 0.0001 |
| | | **Lipid metabolism genes** | | | |
| | | **t1** | | | |
| **SAV2944**[a,b] | *fabB2* | 3-oxoacyl-ACP synthase II | -0.84 | 0.3338 | **0.0410** |
| **SAV217** | *fabC4* | acyl carrier protein | **1.68** | **0.0011** | 0.0000 |
| **SAV492** | *echA1* | enoyl-CoA hydratase | **1.17** | **0.0474** | 0.0014 |

| | | | | | |
|---|---|---|---|---|---|
| **SAV1485**[a] | *plsC1* | 1-acylglycerol-3-phosphate O-acyltransferase | 0.88 | 0.3542 | **0.0471** |
| **SAV1912** | *meaA1* | methylmalonyl-CoA mutase, coenzyme B12-dependent alpha subunit | **1.93** | **0.0003** | 0.0000 |
| **SAV4208** | *ltp3* | nonspecific lipid-transfer protein | **1.16** | **0.0092** | 0.0001 |
| **SAV4209** | | MaoC-like dehydratase | **1.65** | **0.0067** | 0.0001 |
| **SAV4210** | *fadE28* | acyl-CoA dehydrogenase | **1.60** | **0.0035** | 0.0000 |
| | | **t2** | | | |
| **SAV217** | *fabC4* | acyl carrier protein | **1.50** | **0.0048** | 0.0000 |
| **SAV1485**[a] | *plsC1* | 1-acylglycerol-3-phosphate O-acyltransferase | 0.89 | 0.3609 | **0.0463** |
| **SAV2944**[a,b] | *fabB2* | 3-oxoacyl-ACP synthase II | **1.12** | 0.1908 | **0.0098** |
| **SAV3866** | *accA2* | acetyl/propionyl CoA carboxylase alpha subunit | **1.05** | **0.0138** | 0.0001 |
| | | **Energy production genes** | | | |
| | | **t1** | | | |
| **SAV4846** | *nuoJ1* | NADH dehydrogenase I chain J (complex I) | **-1.71** | **0.0434** | 0.0012 |
| **SAV4847** | *nuoK1* | NADH dehydrogenase I chain K (complex I) | **-1.67** | **0.0100** | 0.0002 |
| | | **t2** | | | |
| **SAV2885** | *atpF* | F-type proton-transporting ATPase b chain | **-1.37** | **0.0110** | 0.0001 |
| | | **Transport and external signals processing** | | | |
| | | **t1** | | | |
| **SAV2246** | | simple sugar ABC transporter ATP-binding protein | **-1.10** | **0.0155** | 0.0003 |
| **SAV2247** | | simple sugar ABC transporter substrate-binding protein | **-1.64** | **0.0327** | 0.0009 |
| **SAV2455** | | MFS transporter protein | **-2.42** | **0.0090** | 0.0001 |
| **SAV2609** | | ABC transporter substrate-binding protein | **-1.20** | **0.0229** | 0.0005 |
| **SAV2963** | | ABC transporter permease protein | -0.81 | **0.0396** | 0.0011 |
| **SAV3618** | | transmembrane transport protein | **-1.13** | **0.0314** | 0.0008 |
| **SAV4247** | | ABC transporter permease protein | -0.74 | **0.0252** | 0.0006 |
| **SAV4248**[c] | | ABC transporter ATP-binding protein | **-1.05** | 0.0564 | **0.0018** |
| **SAV4249**[c] | | ABC transporter permease protein | -0.94 | 0.1647 | **0.0099** |
| **SAV4250**[c] | | ABC transporter ATP-binding protein | **-1.39** | 0.0816 | **0.0032** |
| **SAV5915** | *phoA* | alkaline phosphatase | **-2.39** | **0.0013** | 0.0000 |

| | | | | | |
|---|---|---|---|---|---|
| **SAV5971** | *phoC* | acid phosphatase | **-2.63** | **0.0354** | 0.0009 |
| **SAV515** | | oxidoreductase, iron-sulfur subunit | **1.65** | **0.0035** | 0.0000 |
| **SAV600** | *fecC1* | ABC transporter iron(III)/siderophore transport system ATP-binding protein | **2.40** | **0.0003** | 0.0000 |
| **SAV602** | *fecB* | ABC transporter iron(III)/siderophore-binding protein | **2.95** | **0.0006** | 0.0000 |
| **SAV610** | | MFS transporter protein | **1.19** | **0.0166** | 0.0004 |
| **SAV694** | | ABC transporter permease protein | **2.18** | **0.0027** | 0.0000 |
| **SAV695** | | ABC transporter ATP-binding protein | **1.79** | **0.0224** | 0.0005 |
| **SAV4600** | | MFS membrane protein (sulfate transport) | **1.12** | **0.0019** | 0.0000 |
| **SAV6941** | | MFS transporter protein | **3.61** | **0.0000** | 0.0000 |
| **t2** | | | | | |
| **SAV1804** | | hypothetical protein | **-1.00** | **0.0486** | 0.0007 |
| **SAV4066** | | ABC transporter ATP-binding protein | -0.87 | **0.0343** | 0.0004 |
| **SAV4106** | | MFS transporter protein | **-1.51** | **0.0468** | 0.0006 |
| **SAV4247** | | ABC transporter permease protein | -0.73 | **0.0500** | 0.0007 |
| **SAV602** | *fecB* | ABC transporter iron(III)/siderophore-binding protein | **2.24** | **0.0094** | 0.0001 |
| **SAV610** | | MFS transporter protein | **1.43** | **0.0122** | 0.0001 |
| **SAV2436** | | sodium:solute symporter | 0.93 | **0.0297** | 0.0003 |
| **SAV2534** | | serine protease | **1.07** | **0.0297** | 0.0003 |
| **Genes involved in cell envelope biosynthesis and morphological differentiation** | | | | | |
| **t1** | | | | | |
| **SAV2505** | *clgR* | DNA-binding protein | -0.89 | **0.0129** | 0.0002 |
| **SAV2600**[a] | *clpC1* | ATP-dependent Clp protease | -0.58 | 0.7027 | 0.2294 |
| **SAV3188** | | protein sporulation related domain-protein | **-1.39** | **0.0456** | 0.0013 |
| **SAV7486** | *sti2* | subtilisin inhibitor | **-1.09** | **0.0298** | 0.0007 |
| **SAV617** | *ctpB* | cation-transporting P-type ATPase | **1.78** | **0.0009** | 0.0000 |
| **SAV2754** | *ampD1* | N-acetylmuramoyl-L-alanine amidase | **1.38** | **0.0095** | 0.0002 |
| **SAV5456** | *mreC* | rod shape-determining protein | 0.93 | **0.0443** | 0.0012 |
| **SAV6937** | *kipA* | antagonist of KipI | **2.57** | **0.0022** | 0.0000 |
| **SAV6938**[ii] | *kipI* | inhibitor of KinA | **1.51** | 0.0593 | **0.0020** |

| | | | | | |
|---|---|---|---|---|---|
| **t2** | | | | | |
| **SAV2505** | *clgR* | DNA-binding protein | -0.83 | **0.0375** | 0.0004 |
| **SAV3016** | *wblE* | WhiB-family transcriptional regulator | **-1.12** | **0.0451** | 0.0006 |
| **SAV4130** | *bldC* | MerR-family transcriptional regulator | **-1.88** | **0.0279** | 0.0002 |
| **SAV7486** | *sti2* | subtilisin inhibitor | **-1.74** | **0.0039** | 0.0000 |
| **SAV617** | *ctpB* | cation-transporting P-type ATPase | **1.87** | **0.0017** | 0.0000 |
| **Regulatory genes** | | | | | |
| **t1** | | | | | |
| **SAV1081** | | ROK-family transcriptional regulator | **-1.86** | **0.0035** | 0.0000 |
| **SAV2268** | *avaL2* | TetR-family transcriptional regulator (γ-butyrolactone) | **-3.15** | **0.0078** | 0.0001 |
| **SAV2270** | *avaL1* | TetR-family transcriptional regulator (γ-butyrolactone) | **-2.68** | **0.0035** | 0.0000 |
| **SAV2301** | | regulatory protein | **-3.93** | **0.0252** | 0.0006 |
| **SAV2404** | *cutS* | two-component system sensor kinase | **-1.09** | **0.0035** | 0.0000 |
| **SAV2505** | *clgR* | DNA-binding protein | -0.89 | **0.0129** | 0.0002 |
| **SAV2901**[b] | *olmRII* | LuxR-family transcriptional regulator (oligomycin) | **-1.47** | 0.0712 | **0.0026** |
| **SAV2902** | *olmRI* | LuxR-family transcriptional regulator (oligomycin) | **-1.56** | **0.0006** | 0.0000 |
| **SAV2989** | | MarR-family transcriptional regulator (neopentalenolactone) | **-1.06** | **0.0487** | 0.0015 |
| **SAV4327** | | DNA-binding protein | **-1.19** | **0.0336** | 0.0009 |
| **SAV4398** | | regulatory protein | **-1.99** | **0.0396** | 0.0011 |
| **SAV4903** | | TetR-family transcriptional regulator | **-1.41** | **0.0095** | 0.0002 |
| **SAV218** | | transcriptional regulatory protein | **1.11** | **0.0396** | 0.0011 |
| **SAV244** | | MerR-family transcriptional regulator | **2.68** | **0.0029** | 0.0000 |
| **SAV323** | | MerR-family transcriptional regulator | **1.98** | **0.0028** | 0.0000 |
| **SAV343** | | regulatory protein | **1.61** | **0.0148** | 0.0003 |
| **SAV410** | *pteR* | SARP-LAL transcriptional regulator (filipin) | **2.18** | **0.0009** | 0.0000 |
| **SAV448** | | TetR-family transcriptional regulator | **1.12** | **0.0126** | 0.0002 |
| **SAV513** | *bdpJ* | AraC-family transcriptional regulator | **1.45** | **0.0137** | 0.0003 |
| **SAV576** | | TetR-family transcriptional regulator | **2.08** | **0.0014** | 0.0000 |
| **SAV678** | | transcriptional regulator | **1.56** | **0.0045** | 0.0001 |
| **SAV705** | | TetR-family transcriptional regulator | **1.16** | **0.0145** | 0.0003 |

| | | | | | |
|---|---|---|---|---|---|
| **SAV742** | | AraC-family transcriptional regulator | **1.07** | **0.0443** | 0.0012 |
| **SAV831** | | LacI-family transcriptional regulator | **1.21** | **0.0035** | 0.0000 |
| **SAV935** | *aveR* | LuxR-family transcriptional regulator (avermectin) | **2.00** | **0.0049** | 0.0001 |
| **SAV980** | | GntR-family transcriptional regulator | **1.00** | **0.0116** | 0.0000 |
| **SAV5755** | | regulatory protein | **1.62** | **0.0280** | 0.0007 |
| | | **t2** | | | |
| **SAV2505** | *clgR* | DNA-binding protein | -0.83 | **0.0375** | 0.0004 |
| **SAV3705**[iii] | *avaR1* | gamma-butyrolactone receptor protein (avenolide) | -0.92 | 0.1344 | **0.0043** |
| **SAV4130** | *bldC* | MerR-family transcriptional regulator | **-1.88** | **0.0279** | 0.0002 |
| **SAV218** | | transcriptional regulatory protein | **1.22** | **0.0386** | 0.0005 |
| **SAV244** | | MerR-family transcriptional regulator | **2.58** | **0.0079** | 0.0000 |
| **SAV323** | | MerR-family transcriptional regulator | **1.85** | **0.0093** | 0.0000 |
| **SAV410** | *pteR* | SARP-LAL transcriptional regulator (filipin) | **1.44** | **0.0266** | 0.0002 |
| **SAV576** | | TetR-family transcriptional regulator | **1.31** | **0.0436** | 0.0006 |
| **SAV678** | | transcriptional regulator | **1.26** | **0.0343** | 0.0004 |
| **SAV3850** | | transcriptional regulator | **1.41** | **0.0094** | 0.0000 |
| **SAV5754** | | DNA-binding protein | **1.83** | **0.0032** | 0.0000 |
| **SAV5755** | | regulatory protein | **1.83** | **0.0279** | 0.0002 |
| | | **Secondary metabolite genes** | | | |
| | | **t1** | | | |
| **SAV407** | *pteH* | thioesterase | 0.95 | 0.1395 | **0.0076** |
| **SAV408** | *pteG* | cholesterol oxidase | **2.05** | **0.0025** | 0.0000 |
| **SAV411**[b] | *pteE* | ferredoxin (filipin) | **-1.10** | 0.2533 | **0.0233** |
| **SAV412** | *pteD* | cytochrome P450 monooxygenase (filipin) | **-2.74** | **0.0005** | 0.0000 |
| **SAV413** | *pteC* | cytochrome P450 monooxygenase (filipin) | **-2.33** | **0.0004** | 0.0000 |
| **SAV414** | *pteB* | dehydrogenase (filipin) | **-2.34** | **0.0000** | 0.0000 |
| **SAV415**[b] | *pteA5* | modular polyketide synthase (filipin) | **-1.01** | 0.1136 | **0.0054** |
| **SAV416** | *pteA4* | modular polyketide synthase (filipin) | **-1.28** | **0.0095** | 0.0002 |
| **SAV417**[b] | *pteA3* | modular polyketide synthase (filipin) | -0.64 | 0.3514 | **0.0457** |
| **SAV418**[b] | *pteA2* | modular polyketide synthase (filipin) | -0.87 | 0.3639 | 0.0506 |

| | | | | | |
|---|---|---|---|---|---|
| **SAV419** [b] | *pteA1* | modular polyketide synthase (filipin) | -0.80 | 0.4388 | 0.0752 |
| **SAV1136** | *melC1* | tyrosinase co-factor protein | **-1.68** | 0.0776 | **0.0015** |
| **SAV1137**[c] | *melC2* | tyrosinase (melanin) | -0.99 | 0.1151 | **0.0055** |
| **SAV2266**[c] | *avaC* | phosphatase (γ-butyrolactone) | -0.98 | 0.0794 | **0.0030** |
| **SAV2267**[c] | *avaB* | oxidoreductase (γ-butyrolactone) | **-1.20** | 0.2777 | **0.0279** |
| **SAV2269**[c] | *avaA* | gamma-butyrolactone biosynthesis protein (γ- butyrolactone) | **-1.59** | 0.1456 | **0.0083** |
| **SAV2272**[c] | | hypothetical protein (pk3) | **-2.45** | 0.2000 | **0.0150** |
| **SAV2273** | | isomerase (pk3) | **-7.24** | **0.0208** | 0.0005 |
| **SAV2274** | | secreted protein (pk3) | **-5.02** | **0.0473** | 0.0014 |
| **SAV2275** [c] | | transmembrane efflux protein (pk3) | **-4.32** | 0.1124 | **0.0053** |
| **SAV2276**[c] | *fabH7* | 3-oxoacyl-ACP synthase III (pk3) | **-4.24** | 0.1242 | **0.0063** |
| **SAV2277**[c] | | thioesterase (pk3) | **-3.62** | 0.2222 | **0.0183** |
| **SAV2278**[c] | | F420-dependent dehydrogenase (pk3) | **-3.52** | 0.3037 | **0.0327** |
| **SAV2279**[c] | | acyl-CoA synthetase (pk3) | **-2.97** | 0.3522 | **0.0462** |
| **SAV2285**[c] | *rocD3* | ornithine aminotransferase )pks3) | **-3.27** | 0.3205 | **0.0377** |
| **SAV2290**[c] | *fabH3* | 3-oxoacyl-ACP synthase III (pk3) | **-3.30** | 0.3281 | **0.0397** |
| **SAV2724** | *putA* | proline dehydrogenase | **-1.41** | **0.0095** | 0.0002 |
| **SAV2890**[b] | *ccrA1* | crotonyl-CoA reductase (oligomycin) | -0.42 | 0.4368 | 0.0743 |
| **SAV2891**[c] | | hypothetical (oligomycin) | **-1.18** | 0.2280 | **0.0193** |
| **SAV2892**[b] | *olmA4* | modular polyketide synthase (oligomycin) | -0.33 | 0.4760 | 0.0913 |
| **SAV2893** | *olmA5* | modular polyketide synthase (oligomycin) | -0.22 | 0.6421 | 0.1789 |
| **SAV2894**[c] | *olmB* | cytochrome P450 monooxygenase (oligomycin) | **-1.09** | 0.1512 | **0.0087** |
| **SAV2895**[c] | *olmA7* | modular polyketide synthase (oligomycin) | -0.64 | 0.0746 | **0.0028** |
| **SAV2896**[c] | *olmA6* | modular polyketide synthase (oligomycin) | -0.74 | 0.1443 | **0.0080** |
| **SAV2897**[c] | *olmA3* | modular polyketide synthase (oligomycin) | -0.32 | 0.3610 | **0.0493** |
| **SAV2898**[a,b] | *olmA2* | modular polyketide synthase (oligomycin) | -0.31 | 0.5785 | 0.1377 |
| **SAV2899**[a,b] | *olmA1* | modular polyketide synthase (oligomycin) | -0.61 | 0.1817 | **0.0121** |
| **SAV2900**[c] | | P450-like protein (oligomycin) | -0.57 | 0.2474 | **0.0224** |
| **SAV2901** | *olmRII* | LuxR-family transcriptional regulator | **-1.47** | 0.0712 | **0.0026** |
| **SAV2902** | *olmRI* | LuxR-family transcriptional regulator | **-1.56** | **0.0006** | 0.0000 |

| | | | | | |
|---|---|---|---|---|---|
| **SAV2903[c]** | *olmC* | thioesterase (oligomycin) | **-1.86** | 0.1235 | **0.0062** |
| **SAV2990[c]** | *gap1* | glyceraldehyde-3-phosphate dehydrogenase (neopentalenolactone) | **-1.36** | 0.2469 | **0.0222** |
| **SAV2991[c]** | *ptlH* | 1-deoxypentalenic acid 11-beta hydroxylase (neopentalenolactone) | -0.93 | 0.1529 | **0.0089** |
| **SAV2993[c]** | *ptlF* | 1-deoxy-11beta-hydroxypentalenic acid dehydrogenase (neopentalenolactone) | -0.76 | 0.2639 | **0.0251** |
| **SAV2994** | *ptlE* | Baeyer-Villiger monooxygenase (neopentalenolactone) | **-1.30** | **0.0087** | 0.0001 |
| **SAV2995[c]** | *ptlD* | dioxygenase (neopentalenolactone) | **-1.21** | 0.0962 | **0.0041** |
| **SAV3155[c]** | | MbtH-like protein (nrp3) | **-1.18** | 0.2696 | **0.0261** |
| **SAV3156[c]** | *nrps3-1* | non-ribosomal peptide synthetase (nrp3) | **-1.56** | 0.0867 | **0.0035** |
| **SAV3161[c]** | *dapF2* | diaminopimelate epimerase (nrp3) | **-1.03** | 0.5045 | **0.1029** |
| **SAV3162[c]** | | hypothetical protein (nrp3) | **-1.52** | 0.1668 | **0.0101** |
| **SAV3163[c]** | | hypothetical protein (nrp3) | **-1.24** | 0.3227 | **0.0384** |
| **SAV3164** | | hypothetical protein (nrp3) | **-1.59** | **0.0003** | 0.0000 |
| **SAV7112** | *rocD2* | ornithine aminotransferase | **-1.59** | **0.0212** | 0.0005 |
| **SAV7184[c]** | *pks4* | modular polyketide synthase (pks4) | **-3.22** | 0.2980 | **0.0315** |
| **SAV7185[c]** | | UDP-glucose:sterol glucosyltransferase (pks4) | **-3.05** | 0.2753 | **0.0271** |
| **SAV7186[c]** | *cyp26* | cytochrome P450 monooxygenase (pks4) | **-2.88** | 0.2419 | **0.0213** |
| **SAV7469** | *cyp28* | cytochrome P450 monooxygenase | **-1.55** | **0.0359** | 0.0010 |
| **SAV109** | *cyp1* | cytochrome P450 monooxygenase | 0.99 | **0.0319** | 0.0008 |
| **SAV407[b]** | *pteH* | thioesterase (filipin) | 0.95 | 0.1395 | **0.0076** |
| **SAV408** | *pteG* | cholesterol oxidase (filipin) | **2.05** | **0.0025** | 0.0000 |
| **SAV442** | | monooxygenase involved in antibiotic biosynthesis | **1.92** | **0.0116** | 0.0002 |
| **SAV600** | *fecC1* | ABC transporter iron(III)/siderophore transport system ATP-binding protein (nrp6) | **2.40** | **0.0003** | 0.0000 |
| **SAV602** | *fecB* | ABC transporter iron(III)/siderophore-binding protein (nrp6) | **2.95** | **0.0006** | 0.0000 |
| **SAV603** | *nrps6* | non-ribosomal peptide synthetase (nrp6) | 0.68 | 0.3119 | **0.0342** |
| **SAV604** | | hypothetical protein (nrp6) | **1.18** | **0.0224** | 0.0005 |
| **SAV605** | *fadD2* | acyl-CoA synthetase, long-chain fatty acid:CoA ligase (nrp6) | **1.20** | **0.0049** | 0.0001 |
| **SAV606[c]** | | hypothetical protein (nrp6) | 0.72 | 0.3525 | **0.0464** |
| **SAV607** | | taurine catabolism dioxygenase (nrp6) | **1.15** | **0.0017** | 0.0000 |
| **SAV608[c]** | *fabC2* | acyl carrier protein (nrp6) | **1.02** | 0.1136 | **0.0054** |

| | | | | | |
|---|---|---|---|---|---|
| **SAV609**[c] | *fabH4* | 3-oxoacyl-ACP synthase III (nrp6) | **1.15** | 0.1182 | **0.0058** |
| **SAV610** | | MFS transporter protein (nrp6) | **1.19** | **0.0166** | 0.0004 |
| **SAV611** | | beta-hydroxylase (nrp6) | **1.64** | **0.0005** | 0.0000 |
| **SAV803** | *pgmA* | phosphoglucomutase | **1.67** | **0.0155** | 0.0003 |
| **SAV885** | | monooxygenase involved in antibiotic biosynthesis | **2.47** | **0.0003** | 0.0000 |
| **SAV938**[c] | *aveA1* | type I polyketide synthase (avermectin) | **1.18** | 0.1820 | **0.0122** |
| **SAV941**[c] | *aveE* | cytochrome P450 monooxygenase (avermectin) | **1.04** | 0.1376 | **0.0074** |
| **SAV947**[c] | *aveBIII* | glucose-1-phosphate thymidyltransferase (avermectin) | **1.08** | 0.2027 | **0.0154** |
| **SAV951** | *aveBVII* | dTDP-6-deoxy-L-hexose 3-O-methyltransferase (avermectin) | **2.01** | **0.0020** | 0.0000 |
| **SAV952** | *aveBVIII* | dTDP-4-keto-6-deoxy-L-hexose 2,3-reductase (avermectin) | **1.83** | **0.0049** | 0.0001 |
| **SAV953** | *aveG* | thioesterase (avermectin) | **2.20** | **0.0018** | 0.0000 |
| **SAV1002** | | polyketide cyclase | **1.07** | **0.0239** | 0.0006 |
| **SAV2718** | *leuB* | 3-isopropylmalate dehydrogenase | **2.23** | **0.0252** | 0.0006 |
| **SAV3031** | *cyp14* | epi-isozizaene hydroxylase (cytochrome P450) (ezs) | **1.00** | 0.2918 | **0.0301** |
| **SAV3032** | *ezs* | epi-isozizaene synthase (sesquiterpene cyclase) (ezs) | **1.10** | 0.1621 | **0.0097** |
| **SAV3445** | *purN* | phosphoribosylglycinamide formyltransferase | **1.29** | **0.0005** | 0.0000 |
| **SAV3859** | *citA2* | citrate synthase | **1.38** | **0.0144** | 0.0003 |
| **SAV4006** | *mshA* | glycosyltransferase | 0.76 | **0.0422** | 0.0012 |
| **SAV6068** | | secreted cholesterol oxidase | **1.18** | **0.0060** | 0.0001 |
| **SAV7319** | *fhuF2* | ferric iron reductase | **1.26** | **0.0071** | 0.0001 |
| **SAV7320** | *avsA* | siderophore synthetase component (polyhydroxycarboxylate siderophore) | **1.11** | **0.0295** | 0.0007 |
| **SAV7321**[c] | *avsB* | siderophore synthetase component (polyhydroxycarboxylate siderophore) | 0.77 | 0.2474 | **0.0224** |
| **SAV7586** | *mcjB1* | hypothetical protein (micromicin) | **1.16** | **0.0063** | 0.0001 |
| | | **t2** | | | |
| **SAV1136**[c] | *melC1* | tyrosinase co-factor protein (melanin) | **-1.68** | 0.0776 | **0.0015** |
| **SAV1137** | *melC2* | tyrosinase (melanin) | **-1.85** | **0.0078** | 0.0000 |
| **SAV3164** | | hypothetical protein (nrp3) | **-1.43** | **0.0015** | 0.0000 |
| **SAV604** | | hypothetical protein (nrp6) | **1.22** | **0.0357** | 0.0004 |
| **SAV605** | *fadD2* | acyl-CoA synthetase, long-chain fatty acid:CoA ligase (nrp6) | **1.60** | **0.0019** | 0.0000 |

| | | | | | |
|---|---|---|---|---|---|
| **SAV606** | | hypothetical protein (nrp6) | **1.44** | **0.0484** | 0.0007 |
| **SAV607** | | taurine catabolism dioxygenase (nrp6) | **1.75** | **0.0006** | 0.0000 |
| **SAV608[c]** | *fabC2* | acyl carrier protein (nrp6) | 0.99 | 0.1573 | **0.0066** |
| **SAV609[c]** | *fabH4* | 3-oxoacyl-ACP synthase III (nrp6) | **1.08** | 0.1774 | **0.0085** |
| **SAV7586** | *mcjB1* | hypothetical protein (micromicin) | **1.02** | **0.0286** | 0.0003 |
| | | **Miscellaneous** | | | |
| | | **t1** | | | |
| **SAV1149** | | secreted esterase | **-0.86** | **0.0212** | 0.0005 |
| **SAV1745** | | monooxygenase | **-1.01** | **0.0116** | 0.0002 |
| **SAV2622** | | secreted protein | **-1.83** | **0.0020** | 0.0000 |
| **SAV2764** | | secreted metallopeptidase | **-1.35** | **0.0320** | 0.0008 |
| **SAV3319** | | SAM-P45 peptidase | **-2.35** | **0.0022** | 0.0000 |
| **SAV4202** | | secreted protein | **-1.19** | **0.0208** | 0.0005 |
| **SAV4718** | | hypothetical protein | **-1.05** | **0.0321** | 0.0008 |
| **SAV5292** | | secreted protein | **-1.37** | **0.0049** | 0.0001 |
| **SAV5827** | | secreted protein | **-2.44** | **0.0035** | 0.0000 |
| **SAV6208** | | trypsin-like protease | -0.82 | **0.0474** | 0.0014 |
| **SAV6295** | | carboxypeptidase T (secreted zinc-binding carbopeptidase) | **-2.40** | **0.0438** | 0.0012 |
| **SAV194** | | acetyltransferase | **1.92** | **0.0001** | 0.0000 |
| **SAV200** | | large secreted protein | **1.35** | **0.0487** | 0.0015 |
| **SAV468** | | oxygenase | **1.92** | **0.0468** | 0.0014 |
| **SAV613** | | secreted protein | **1.76** | **0.0024** | 0.0000 |
| **SAV615** | | secreted protein | **2.07** | **0.0033** | 0.0000 |
| **SAV616** | | secreted protein | **1.81** | **0.0047** | 0.0001 |
| **SAV769** | | dioxygenase | **2.38** | **0.0003** | 0.0000 |
| **SAV787** | | choramphenicol phosphotransferase | **1.35** | **0.0038** | 0.0000 |
| **SAV871** | | Fe(II)/alpha-ketoglutarate dependent hydroxylase | **1.27** | **0.0088** | 0.0001 |
| **SAV897** | | secreted alpha-amylase inhibitor | **2.23** | **0.0343** | 0.0009 |
| **SAV899** | | secreted protein | **2.84** | **0.0001** | 0.0000 |
| **SAV903** | | membrane protein | **1.10** | **0.0110** | 0.0002 |

| | | | | |
|---|---|---|---|---|
| **SAV923** | hydrolase | **1.08** | **0.0156** | 0.0003 |
| **SAV974** | non-heme chloroperoxidase | **1.31** | **0.0224** | 0.0005 |
| **SAV990** | secreted protein | **1.81** | **0.0022** | 0.0001 |
| **SAV993** | cysteine transferase | **1.64** | **0.0077** | 0.0001 |
| **SAV996** | secreted protein | **1.09** | **0.0027** | 0.0000 |
| **SAV1008** | secreted protein | **2.54** | **0.0002** | 0.0000 |
| **SAV1048** | DNA-binding protein | **1.52** | **0.0070** | 0.0001 |
| **SAV1749** | phosphoesterase (SimX4 homolog) | **1.15** | **0.0125** | 0.0002 |
| **SAV2164** | hydrolase | 0.92 | **0.0429** | 0.0012 |
| **SAV4008** | NLP/P60-family secreted protein (PgpA peptidase) | 0.78 | **0.0449** | 0.0013 |
| **SAV4330** | membrane protein | **1.67** | **0.0088** | 0.0001 |
| **SAV4668** | secreted protein | 0.76 | **0.0456** | 0.0013 |
| **SAV5004** | secreted beta-lactamase | **1.18** | **0.0437** | 0.0014 |
| **SAV6908** | membrane protein | 0.79 | **0.0496** | 0.0015 |
| **SAV6939** | lactam utilization protein | **2.21** | **0.0035** | 0.0000 |
| **SAV7350** | integral membrane protein | **2.96** | **0.0030** | 0.0000 |
| | | **t2** | | |
| **SAV3095** | ATP-binding protein | **-1.31** | **0.0372** | 0.0004 |
| **SAV51** | acetyltransferase | **1.19** | **0.0372** | 0.0004 |
| **SAV194** | acetyltransferase | 0.91 | **0.0321** | 0.0003 |
| **SAV615** | secreted protein | **2.51** | **0.0020** | 0.0000 |
| **SAV812** | small hydrophilic protein | 0.68 | **0.0484** | 0.0007 |
| **SAV871** | Fe(II)/alpha-ketoglutarate dependent hydroxylase | **1.18** | **0.0283** | 0.0003 |
| **SAV899** | secreted protein | **2.71** | **0.0006** | 0.0283 |
| **SAV923** | hydrolase | **1.47** | **0.0050** | 0.0000 |
| **SAV990** | secreted protein | **1.47** | **0.0156** | 0.0001 |
| **SAV996** | secreted protein | 0.92 | **0.0162** | 0.0000 |
| **SAV1008** | secreted protein | **1.61** | **0.0085** | 0.0000 |
| **SAV1048** | DNA-binding protein | **1.43** | **0.0204** | 0.0002 |
| **SAV1275** | stress-inducible protein | 0.90 | **0.0484** | 00007 |

| | | | | | |
|---|---|---|---|---|---|
| **SAV1499** | | integral membrane protein | **1.27** | **0.0279** | 0.0002 |
| **SAV1501** | | secreted protein | **1.34** | **0.0455** | 0.0006 |
| **SAV1512** | | secreted protein | **1.07** | **0.0184** | 0.0001 |
| **SAV4438** | | secreted protein | **2.29** | **0.0404** | 0.0005 |
| **SAV6553** | *sprD1* | streptogrisin D (secreted serine protease) | **1.74** | **0.0486** | 0.0001 |
| **SAV7350** | | integral membrane protein | **2.56** | **0.0160** | 0.0001 |

[i] PimM binds to its promoter region (Vicente et al., 2015)
[ii] Gene included considering previous (Vicente et al., 2015) or present RT-qPCR results
[iii] Gene included because its transcription profile matches those of genes functionally related
